# RNA Helicase DDX3 Regulates RAD51 Localization and DNA Damage Repair in Ewing Sarcoma

**DOI:** 10.1101/2023.06.10.544474

**Authors:** Matthew E. Randolph, Marwa Afifi, Aparna Gorthi, Rachel Weil, Breelyn A. Wilky, Joshua Weinreb, Paul Ciero, Natalie ter Hoeve, Paul J. van Diest, Venu Raman, Alexander J. R. Bishop, David M. Loeb

**Author notes:** **Corresponding Author:** David M. Loeb, MD, PhD; Albert Einstein College of Medicine, 1300 Morris Park Ave., Van Etten 6A-04D, Bronx, NY 10461.

## Abstract

We previously demonstrated that RNA helicase DDX3X (DDX3) can be a therapeutic target in Ewing sarcoma (EWS), but its role in EWS biology remains unclear. The present work demonstrates that DDX3 plays a unique role in DNA damage repair (DDR). We show that DDX3 interacts with several proteins involved in homologous recombination, including RAD51, RECQL1, RPA32, and XRCC2. In particular, DDX3 colocalizes with RAD51 and RNA:DNA hybrid structures in the cytoplasm of EWS cells. Inhibition of DDX3 RNA helicase activity increases cytoplasmic RNA:DNA hybrids, sequestering RAD51 in the cytoplasm, which impairs nuclear translocation of RAD51 to sites of double-stranded DNA breaks thus increasing sensitivity of EWS to radiation treatment, both *in vitro* and *in vivo*. This discovery lays the foundation for exploring new therapeutic approaches directed at manipulating DDR protein localization in solid tumors.

## Introduction

Ewing sarcoma (EWS) is the second most common high-grade bone sarcoma in children and adolescents. Overall survival of EWS patients is less than 30% for patients with metastatic or recurrent disease despite aggressive chemotherapy, radiation and/or surgery ^1, 2^. EWS is characterized by a chromosomal translocation of the EWS RNA binding protein 1 (*EWSR1*) with an erythroblast transformation specific (*ETS*) family gene or ETS-related gene such as friend leukemia integration 1 transcription factor (*FLI-1*) or transcriptional regulator ERG (*ERG*), in 85% and 10% of all EWS cases, respectively ^1^. The resultant fusion protein acts as a driver for the oncologic biology of EWS with few somatic mutations contributing to the phenotype ^3, 4^. To date, no systemic therapies exist that prolong overall survival of children with metastatic or recurrent EWS. New therapeutic targets, or approaches that increase the potency or effectiveness of current therapeutics, are desperately needed.

DEAD/H box RNA helicases are a superfamily of ATPase dependent RNA helicases which have a conserved amino acid sequence (Asp-Glu-Ala-Asp/His). These RNA helicases unwind and remodel RNA:RNA duplexes, RNA:DNA hybrids and messenger ribonucleoprotein complexes ^5, 6^. ATP dependent RNA helicases, such as DDX3X (DDX3), are ubiquitous enzymes involved in multiple facets of RNA metabolism and are increasingly recognized as important contributors to cancer pathogenesis ^7, 8^. DDX3 has been specifically implicated in the pathogenesis of EWS ^9,10^. We have previously demonstrated that multiple sarcoma cell lines, including EWS, and primary sarcoma patient-derived xenograft (PDX) models express high levels of DDX3 ^10^. We have also shown that treatment with RK-33, a small molecule inhibitor of DDX3 RNA helicase activity ^11–13^, is selectively cytotoxic to EWS cell lines, compared with normal mesenchymal cells ^10^. Additionally, RK-33 sensitivity in PDX models correlates with DDX3 expression ^10^, suggesting that interfering with DDX3 function may be a viable treatment strategy for EWS.

To further understand the underlying mechanisms of this phenotype, proteomic analysis of EWS cell lines with either shRNA knockdown or chemical inhibition of DDX3 was performed. These studies revealed several cellular pathways that are affected by DDX3 impairment, one of which is the DNA damage repair (DDR) pathway ^10^. Induction of DNA damage via ionizing radiation (IR) is a treatment modality for a subset of EWS patients with recurrent or metastatic disease ^14^ as well as for patients with unresectable primary tumors ^15^. Therefore, we investigated whether DDX3 contributes to DDR in EWS and if impairment of DDX3 helicase activity with RK-33 in EWS could be leveraged as a radiosensitizing strategy.

## Results

### Expression of DDX3 is prevalent in EWS patients and is negatively associated with long-term survival

Previously, we demonstrated that multiple sarcoma cell lines and primary sarcoma PDX models express high levels of DDX3 ^10^; however, the frequency and significance of DDX3 expression in EWS patient samples had not been assessed. To determine whether DDX3 expression is prevalent in EWS patients, immunohistochemical analysis of DDX3 expression was performed using an EWS tissue microarray. High DDX3 expression was present in the majority of EWSs examined (Figure 1A, 1B). To evaluate the prognostic impact of DDX3 expression in EWS patients, we analyzed a previously reported data set (GEO ID: gse63157) of mRNA transcripts obtained from EWS patients ^16^ using the R2 Genomics Platform (https://r2.amc.nl). Elevated DDX3 mRNA levels correlated with worse event-free (p=0.0014) and overall survival (p=0.044) rates compared to patients whose tumors expressed low levels of DDX3 mRNA (Figure 1C, 1D). This correlation further supports the clinical value of targeting DDX3 function as a potentially beneficial therapy for EWS patients.

**Figure 1.**
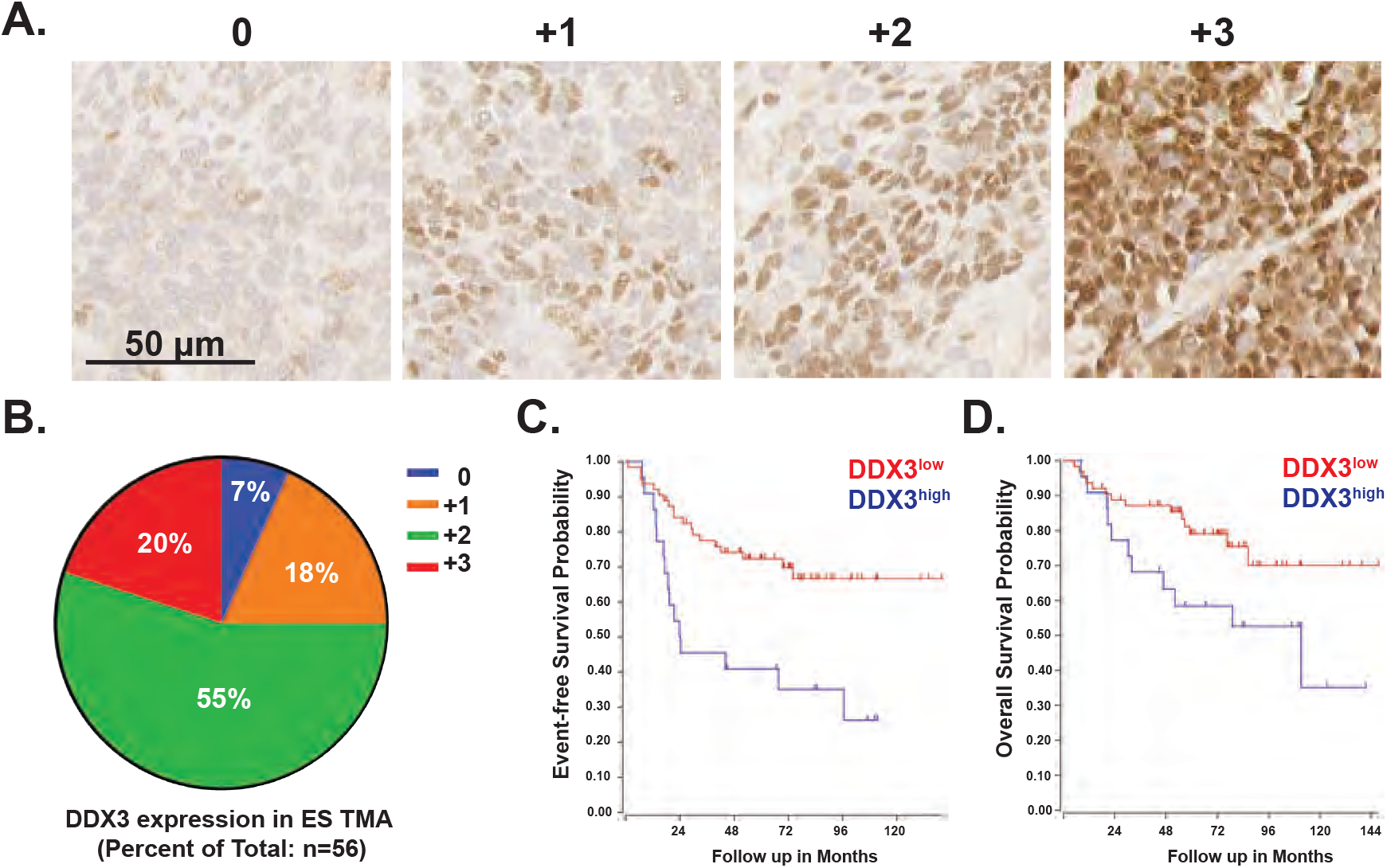
Elevated DDX3 expression in Ewing sarcoma is associated with poor prognosis. (A) Immunohistochemical staining of Ewing sarcoma tissue microarray (TMA). Representative images of DDX3 protein expression. Mag bar = 50 µm. (B) Pie chart showing the distribution of DDX3 expression among TMA samples (n = 56. (C) Kaplan-Meier curves demonstrating event-free survival (EFS) and overall survival (OS) of EWS patients’ tumors with either low (red) (n = 63) or high (blue) (n = 22) levels of DDX3X mRNA transcripts. EFS: raw p = 0.0014 by log rank (Mantel-Cox) test. OS: raw p = 0.044 by log rank (Mantel-Cox) test.

### Double-stranded DDR is abrogated by inhibition of DDX3 helicase activity following IR

To elucidate the underlying contribution of DDX3 function to EWS biology, we previously used a proteomic approach to identify cellular pathways/processes in EWS that were uniformly altered with both genetic and chemical inhibition of DDX3 ^10^. One of the major cellular processes altered with DDX3 inhibition was DDR ^10^. To examine the effects of DDX3 inhibition on DDR, we irradiated (2 Gy) EWS cell lines that either stably expressed shRNA against DDX3 ^10^ or were treated with RK-33. Resultant double-stranded DNA breaks were quantified via the immunofluorescent presence of phosphorylated histone H2A variant H2A.X (γ-H2A.X) foci (Figure 2A) ^17^. To inhibit DDX3 function genetically, we utilized our previously established stable DDX3 knockdown MHH-ES-1 EWS cell lines that constitutively express shDDX3 with a verified knockdown (KD) of 70-80% by Western blot ^10^. In the presence of decreased constitutive levels of DDX3, we observed both higher basal levels of DSBs in KD verses control cell lines and impairment of DDX3^KD^ cells to resolve DSBs by 24 hours post-IR, suggesting DDX3 contributes to the maintenance of genomic integrity in EWS (Figure 2B). Additionally, chemical inhibition of DDX3 RNA helicase activity with RK-33 treatment significantly inhibited the restoration of double-stranded DNA break abundance back to basal levels within 24 hours of IR in three independent EWS cell lines (Figure 2C). While the overall effect was consistent, the magnitude of the inhibition varied among the EWS cell lines, as would be expected in a clinical setting. Importantly, clonogenic assays demonstrated that survival of EWS cells was significantly impaired following the combined treatment of RK-33 with IR (Figure 2D, 2E). Thus, pharmacologic or genetic inhibition of DDX3 attenuates DDR in EWS cells, resulting in increased cell death.

**Figure 2.**
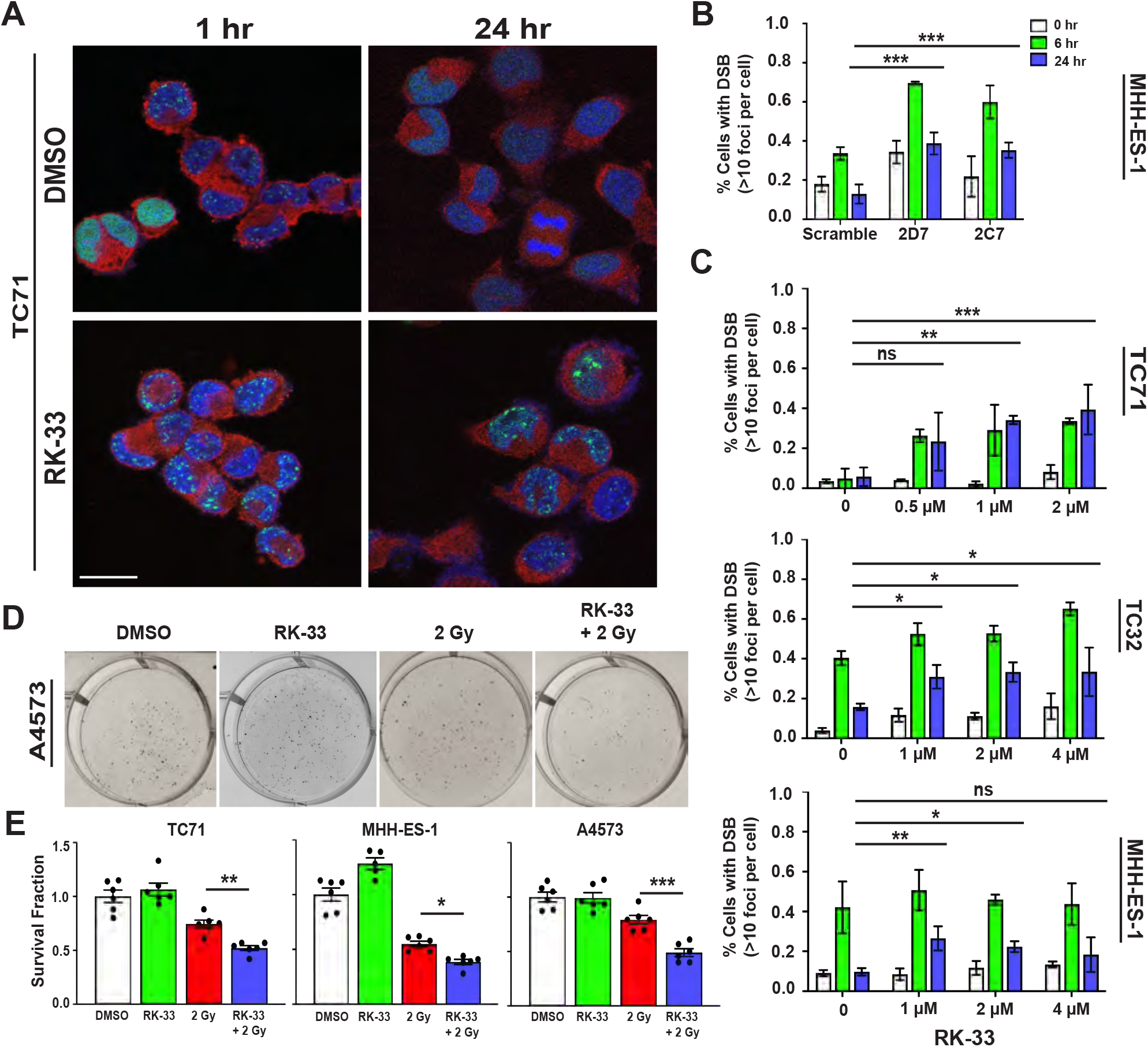
Inhibition of DDX3 radiosensitizes Ewing sarcoma. (A) Immunofluorescent images of TC71 EWS cells at 1 or 24 hours following 2 Gy irradiation that were treated with either DMSO (top) or 2 µM RK-33 (bottom). Green = γ-H2A.X detection, marking double-stranded DNA breaks (DSB); Red = DDX3; Blue = DAPI stain. Mag bar = 20 µm. (B) Quantitation of DSBs in stable genetically modified shDDX3 MHH-ES-1 cell lines 2D7 and 2C7 at 0 (*i.e.* no treatment), 6, and 24 hours following 2 Gy irradiation. Data are representative of three independent experiments. Data are mean ± SD. *p<0.05 and ***p<0.001 determined by Two-Way ANOVA followed by Šídák’s multiple comparisons test. (C) Quantitation of DSBs in three independent Ewing sarcoma cell lines (TC71, MHH-ES-1, and TC32) where cells were irradiated with 2 Gy in the presence of 0, 0.5, 1, 2, or 4 µM RK-33. Data represent three independent experiments per cell line. Data are mean ± SD. *p<0.05 and ***p<0.001 determined by Two-Way ANOVA followed by Šídák’s multiple comparisons test. (D-E) A4573 EWS cells were treated with either DMSO, 2 Gy, 2 µM RK33, or 2 µM RK-33 + 2 Gy and plated at densities of 400 cells/well 6 hours post-irradiation. Cells were then grown in conditioned media for 5 days and stained with crystal violet for (D) visualization and (E) quantification of clonogenic survival fractions. (n = 6 technical replicates per cohort per cell line). Results represent one experiment of three independent experiments per cell line. Data are mean ± SEM. *p<0.05, **p<0.01 and ***p<0.001 determined by One-Way ANOVA followed by Šídák’s multiple comparisons test.

Because inhibition of DDX3 impairs the repair of double-stranded DNA breaks up to 24 hours after IR, we tested whether chemical impairment of DDX3 with RK-33 could serve as a radiosensitizing approach in EWS PDX models. Two EWS PDX models, EWS4 and JHH-ESX-3, that respectively express low (DDX3^low^) and high (DDX3^high^) levels of DDX3 protein (Figure 3A), were utilized to test whether RK-33 could act as a radiosensitizing agent when combined with sub therapeutic doses of columnated IR (10 Gy; Figure 3B). Single agent treatment with a sub-therapeutic dose of 50 mg/kg RK-33 (p=0.627) did not significantly alter tumor volume, while IR, alone (p=0.0229) or in combination with RK-33 (p=0.0021), resulted in a significant decrease of volume in the DDX3^low^ PDX (Figure 3C). Importantly, the DDX3^high^ PDX was completely ablated (p<0.0001) by combined DDX3 inhibition and IR as early as 10 days post-IR (Figure 3D inset), an effect that was maintained for several weeks before tumor recurrence (Figure 3D). Additionally, combination therapy of RK-33 with IR significantly improved the survival rate of mice implanted with DDX3^high^ (p<0.0001) but not DDX3^low^ (p=0.2802) EWS PDXs (Figure 3E, F), consistent with DDX3^high^ tumors having a greater dependency on DDX3, as we have previously reported ^10^. Taken together, these data demonstrate that the impact of RK-33 is dependent upon the level of expression of its target, DDX3, and provide support for the development of RK-33 as a radiosensitizing agent in EWS patients whose tumors demonstrate high levels of DDX3 protein expression.

**Figure 3.**
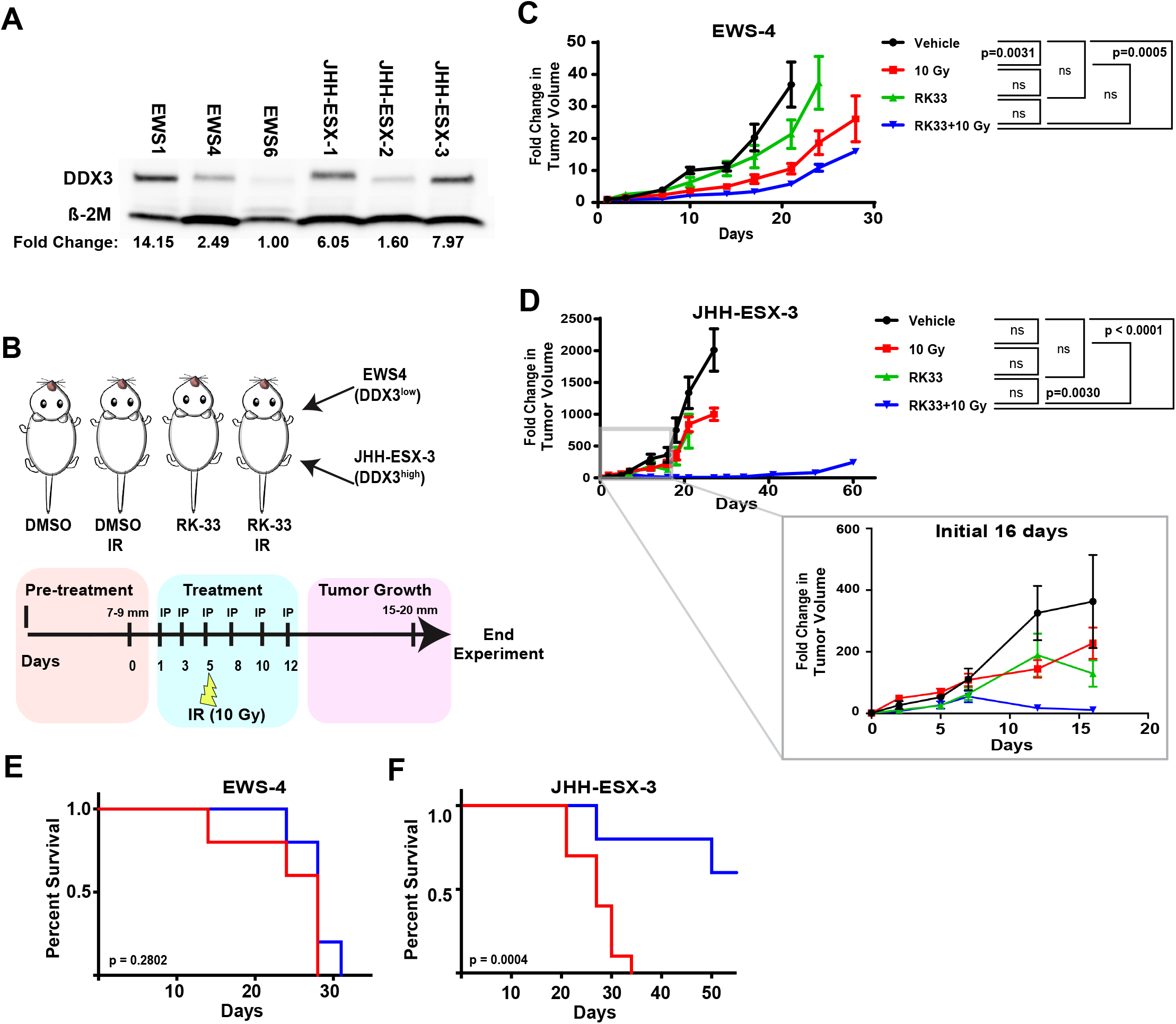
RK-33 induces radiosensitization in Ewing sarcoma xenograft models expressing high levels of DDX3. (A) Quantitation of DDX3 protein expression in six independent Ewing sarcoma PDX models. All samples were normalized using the corresponding loading control, β-2M. Fold changes of protein abundance were calculated by comparing normalized band densities to EWS6 abundance and are noted below each sample for comparison. β-2M = beta 2-microglobulin. (B) Schematic representation of treatment schedule and cohorts of two independent EWS patient-derived xenograft (PDX) NSG mouse models: EWS4 (DDX3^low^) and JHH-ESX-3 (DDX3^high^). Tumor chunks were implanted subcutaneously. Upon reaching a tumor diameter of 7-9 mm, mice received every-other day intraperitoneal (IP) injections of either DMSO or RK-33 (25 mg/kg) for two weeks with radiation cohorts (IR) receiving 10 Gy radiation administered 6 hours following drug treatment on day 5. Mice were euthanized and tumors collected after growing to a diameter of 15-20 mm or Day 60, whichever came first. Tumors were measured with calipers twice weekly. (C, D) Tumor volumes of DDX3^low^ (C) and DDX3^high^ (D) were measured and calculated over a period of 29 to 60 days, respectively. Inset shows changes in DDX3^high^ tumor volume during the first 16 days post-treatment demonstrating tumor ablation of RK-33/10 Gy cohort by day 16. Results represent one experiment of 4-10 mice per cohort per PDX. Data are mean ± SEM. Statistical significance was determined by Two-Way ANOVA followed by Tukey’s multiple comparisons test at Day 21. (E, F) Kaplan-Meier curves demonstrating survival of 10 Gy verses RK-33/10 Gy cohorts of DDX3^low^ (E) and DDX3^high^ (F) PDXs. Statistical significance was determined by log rank (Mantel-Cox) test.

### Homologous and non-homologous DDR pathways are impaired by DDX3 inhibition

Recent studies suggest that EWS inherently has impaired homology-directed DNA damage repair (HR) owing to sequestration of BRCA1 by hyperphosphorylated RNAPII via an EWS-FLI1 dependent mechanism ^18^. Therefore, we hypothesized that RK-33 inhibition of DDX3 in EWS could further exacerbate the tumor’s basal impaired ability to repair genomic damage, resulting in enhanced persistence of double-stranded DNA breaks, thus contributing to the pronounced *in vivo* radiosensitization phenotype. To determine which DDR pathways were affected by DDX3 abrogation, we utilized I-SceI GFP reporter constructs in U2OS cells ^19, 20^ to identify and quantify double-stranded DNA break repair mechanisms including HR and total end joining, which is reflective of non homologous end joining (NHEJ). These reporters consist of two non-functional GFP genes in tandem. The upstream gene has a mutation introducing the I-SceI recognition site and the downstream gene is an internal fragment. Upon induction of an I-SceI double strand break, the downstream gene operates as a template to repair the break when HR or NHEJ is proficient, resulting in functional GFP gene expression that can be measured by flow cytometry. Knockdown of DDX3 (Figure 4A) resulted in a significant decrease in both HR (p<0.001) and NHEJ activity (p<0.001) as measured by this assay (Figure 4B). In contrast, single-strand annealing DNA repair was unaffected with DDX3 impairment (Figure 4B). These data suggest a role for DDX3 in both homologous and non homologous double-stranded DNA break repair.

Repair of double-stranded DNA breaks (DSB), either through HR or NHEJ, involves multiple proteins and signaling cascades which direct the repair pathways ^21^. Given the role of DDX3 in translational regulation ^22–24^, we began to investigate how DDX3 inhibition impairs DDR by assessing changes in protein abundance and signaling following RK-33 treatment of EWS cells with or without IR. TC71 or MHH-ES-1 cells were pre-treated with either DMSO or RK-33 for one hour prior to receiving 2 Gy IR. Cells were collected for protein analysis at either 1- or 24-hours post-IR. Western blots were performed assessing a panel of proteins involved with DSB repair (DSBR) (Figure S1) (n=2-3 experiments per cell line). We did not observe any consistent modulations in the abundance of proteins involved in either the HR or NHEJ repair pathways. Additionally, no significant alterations in DDR signaling, such as the phosphorylation of BRCA1 at Ser1423 ^25, 26^, occurred with RK-33 treatment following IR, nor did we find evidence of RK-33 altering CHK1 Ser345 phosphorylation or replication protein A subunit 32 (RPA32) on Ser4/Ser8, markers for replication fork stress ^27, 28^. These data suggest that DDX3 does not regulate DSBR by grossly altering the abundance or signaling cascades of proteins involved in DSBR or replication stress.

### DDX3 associates with HR proteins but not at sites of double-stranded DNA breaks

We next investigated whether DDX3 might physically interact with proteins of the HR and NHEJ repair pathways. Immunoprecipitations of DDX3 were performed and analyzed via Western blot for a panel of DDR proteins involved in HR and/or NHEJ repair (Figure 4C). Proteins associated with NHEJ repair, such as Ku70 and Ku86 [reviewed in ^29^] did not interact with DDX3, whereas ATP dependent DNA helicase Q1 (RECQL1), RPA32, DNA repair protein RAD51 homolog 1 (RAD51), and X-ray repair cross-complementing 2 (XRCC2) [reviewed in ^21, 30, 31^], all of which contribute to homology-directed repair, co-immunoprecipitated with DDX3 in three independent EWS cell lines (Figure 4C, Figure S2A).

Given the observation that DDX3 interacts with several DNA repair proteins, but these DDR proteins are generally considered to be localized to the nucleus while the majority of DDX3 is cytoplasmic, we evaluated the subcellular localization of these proteins. A detectable amount of DDX3 was noted in the nuclear fraction. However, more interesting was the observation that the DDX3 DDR interacting proteins RPA32, RAD51, XRCC2 and, to a lesser extent, RECQL1, were found not only in the nuclear fraction but also in the cytoplasm (Figure 4D, Figure S2B); eIF4E, H2A.X and NUP205 provide controls for the quality of the fractionation. By immunofluorescence we did not observe any DDX3 colocalization at sites of DSBs, as identified by the presence of γ-H2A.X foci (Figure 4E); the majority of DDX3 was cytoplasmic which also agrees with our immunohistochemical assessment of EWS microarray samples (Figure 1; 96.4% (54/56) majority was cytoplasmic). These data suggest that endogenous DDX3 does not directly contribute to DSBR at the site of DNA damage, but possibly through interactions with DDR proteins in the cytoplasm of EWS cells.

**Figure 4.**
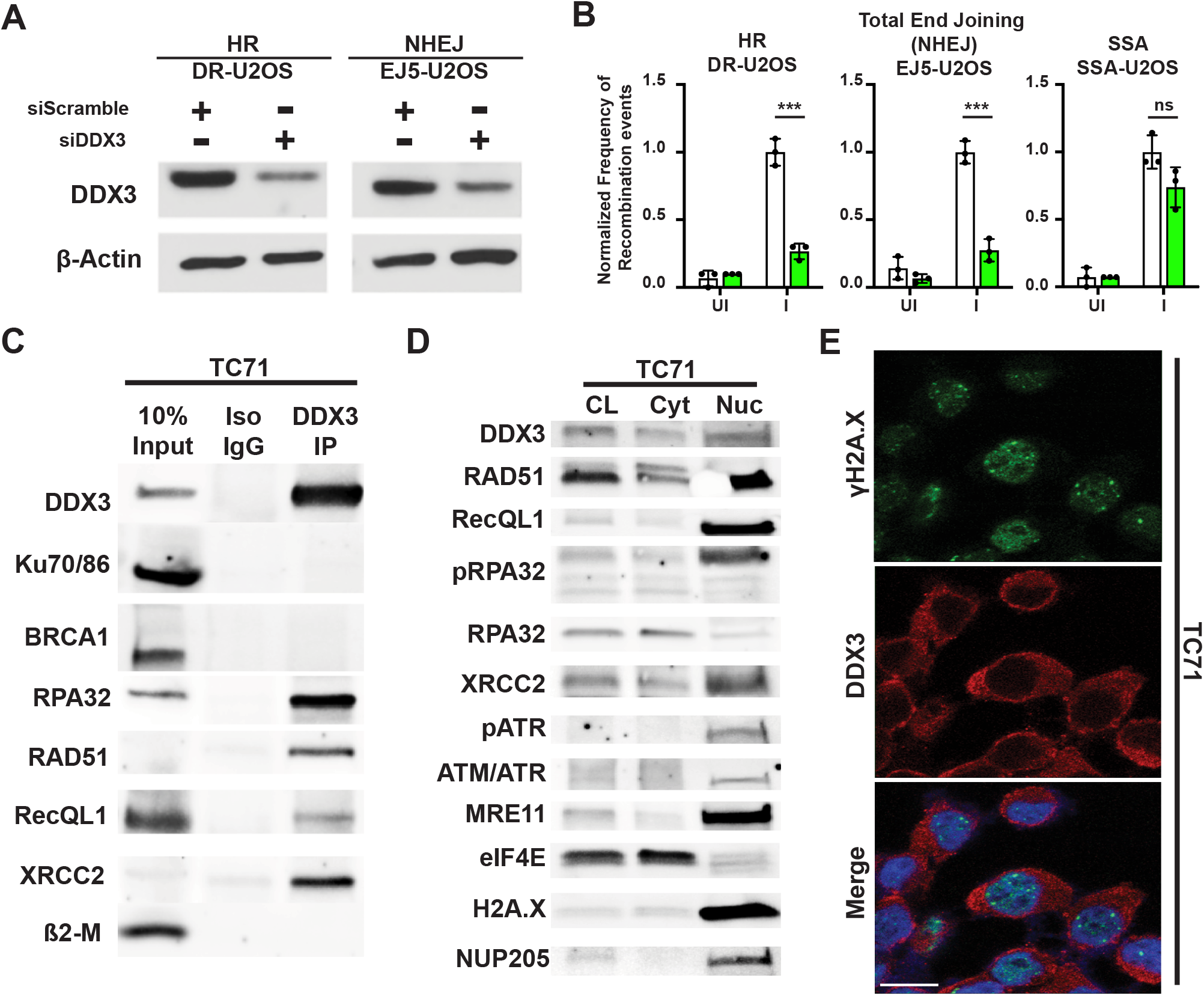
DDX3 interacts with components of the homologous DNA damage repair pathway. (A) Western blots of DR-U2OS and EJ5-U2OS cell lysates following siRNA knockdown of DDX3. (B) DR-U2OS, EJ5-U2OS, and SSA-U2OS cell lines were treated with either scramble or siDDX3 and then transfected with the I*Sce*I-pCAGGS vector or an empty vector to induce DNA damage. Effective DNA damage repair was visualized by induction of GFP expression and quantified using flow cytometry. Results represent three independent experiments per cell line. Data represent frequency of DNA recombination events SEM. ***p<0.001 determined by multiple unpaired t-tests followed by Šídák’s multiple comparisons test. I = induction of DNA damage with I*Sce*I-pCAGGS; UI = non induction with empty pCAGGS vector; HR = homologous recombination; NHEJ = nonhomologous end joining; SSA = single-strand annealing. (C) Immunoprecipitation (IP) of TC71 EWS cell lysates using anti-DDX3 antibodies conjugated to magnetic beads. Western blots demonstrate co immunoprecipitation of various DNA damage repair proteins with endogenous DDX3. Iso IgG = immunoprecipitation of TC71 cell lysates using control isotype antibodies. Results are representative of 3 independent experiments. See also Figure S2. (D) Subcellular fractionations of TC71 cells demonstrate the presence of DDX3, RAD51, RECQL1, RPA32 and XRCC2 in both cytoplasmic and nuclear compartments. Results are representative of 3 independent experiments. See also Figure S2. (E) Immunofluorescent images of TC71 EWS cells 6 hours after treatment with 5 µM RK-33 and 2 Gy irradiation. Green = γ-H2A.X staining of double-stranded DNA breaks (DSB); Red = DDX3; Blue = DAPI stain. Mag bar = 20 µm.

### RNA helicase DDX3 interacts with cytoplasmic oligonucleotide substrates in EWS

Successful DDR relies on regulated translocation of several DDR proteins from the cytoplasm to the nucleus in response to DNA damage, as has been previously investigated with BRCA1, BRCA2, BRCA1-associated RING domain protein 1 (BARD1), TP53-binding protein 1 (53BP1), BRCA1-associated ATM activator 1 (BRAT1), exonuclease 1 (EXO1), XRCC4, and RAD51 ^32–37^. Considering that a) DDX3 localization in EWS is predominantly cytoplasmic, b) endogenous DDX3 does not localize to sites of DSBs, and c) DDX3 interacts with HR proteins, we hypothesized that impairment of DDX3 helicase activity could sequester DDX3, and potentially DDR proteins, in the cytoplasm, thus preventing appropriate translocation into the nucleus, resulting in an inhibition of DDR.

Recent studies have demonstrated replication stress in EWS that appears to be driven by the EWS-FLI1 fusion protein ^18, 38, 39^. More specifically, EWS-FLI1 expression causes hyperphosphorylation of RNAPII, increasing transcription and resulting in increased R-loop abundance, replication stress and DNA damage ^18^. High levels of DNA replication stress have also been associated with cytoplasmic accumulation of genomic DNA [reviewed in ^40^]. Based on this concept we looked for cytoplasmic DNA and noted its presence as well as micronuclei in multiple EWS cell lines, EWS PDXs, and EWS patient samples (Figure 5A-C). Additionally, DDX3 has been shown to interact with multiple oligonucleotide substrates *in vitro*, including RNA:RNA duplexes, RNA:DNA hybrids, and single-stranded DNA, with decreasing affinity respectively ^22, 41^. Given this, we looked to determine if these different forms of nucleic acid structures were present in the cytoplasm of EWS cells. Intriguingly, we discovered that both single-stranded DNA (ssDNA) and RNA:DNA hybrid structures are present in the cytoplasm of EWS cells with the majority of cytoplasmic ssDNA substrates colocalizing with RNA:DNA hybrids (Figure 5D). Indeed, cytoplasmic S9.6 staining was consistently observed in EWS but not in primary umbilical chord-derived mesenchymal stem cells (Figure 5E). We confirmed the identity of the cytoplasmic RNA:DNA hybrid structures by overexpressing RNaseH1, an *in vitro* technique commonly used to resolve nuclear RNA:DNA hybrid structures, such as transcriptional R-loops, by specifically degrading the RNA strand ^42–44^. Lentivirus driven overexpression of RNaseH1^WT^ in EWS cell lines significantly reduced cytoplasmic RNA:DNA hybrid abundance, as demonstrated by S9.6 antibody staining ^45^, compared to the enzymatically inactive mutant RNaseH1^D210N 46^ (Figure 5F,5G). Conversely, cytoplasmic RNA:RNA duplexes, identified with the J2 antibody specific for dsRNA ^47^, increased following RNaseH1^WT^ overexpression (Figure 5H), which is consistent with dissociated RNA from RNA:DNA hybrids forming secondary and tertiary structures. Importantly, cytoplasmic J2 staining was minimal in EWS overexpressing the RNaseH1^D210N^ control (Figure 5H), suggesting that dsRNA is less likely the nucleic acid substrate responsible for the cytoplasmic localization of DDX3. Next, we examined whether DDX3 colocalized with cytoplasmic RNA:DNA hybrids or with ssDNA. Although DDX3 colocalized with both substrates, DDX3 demonstrated a more robust colocalization with cytoplasmic RNA:DNA hybrids (Figure 5I) compared to ssDNA under basal conditions (Figure S3). Together, these findings suggest that DDX3 preferentially interacts with cytoplasmic RNA:DNA oligonucleotide substrates in EWS, which could account for the cytoplasmic localization of DDX3.

**Figure 5.**
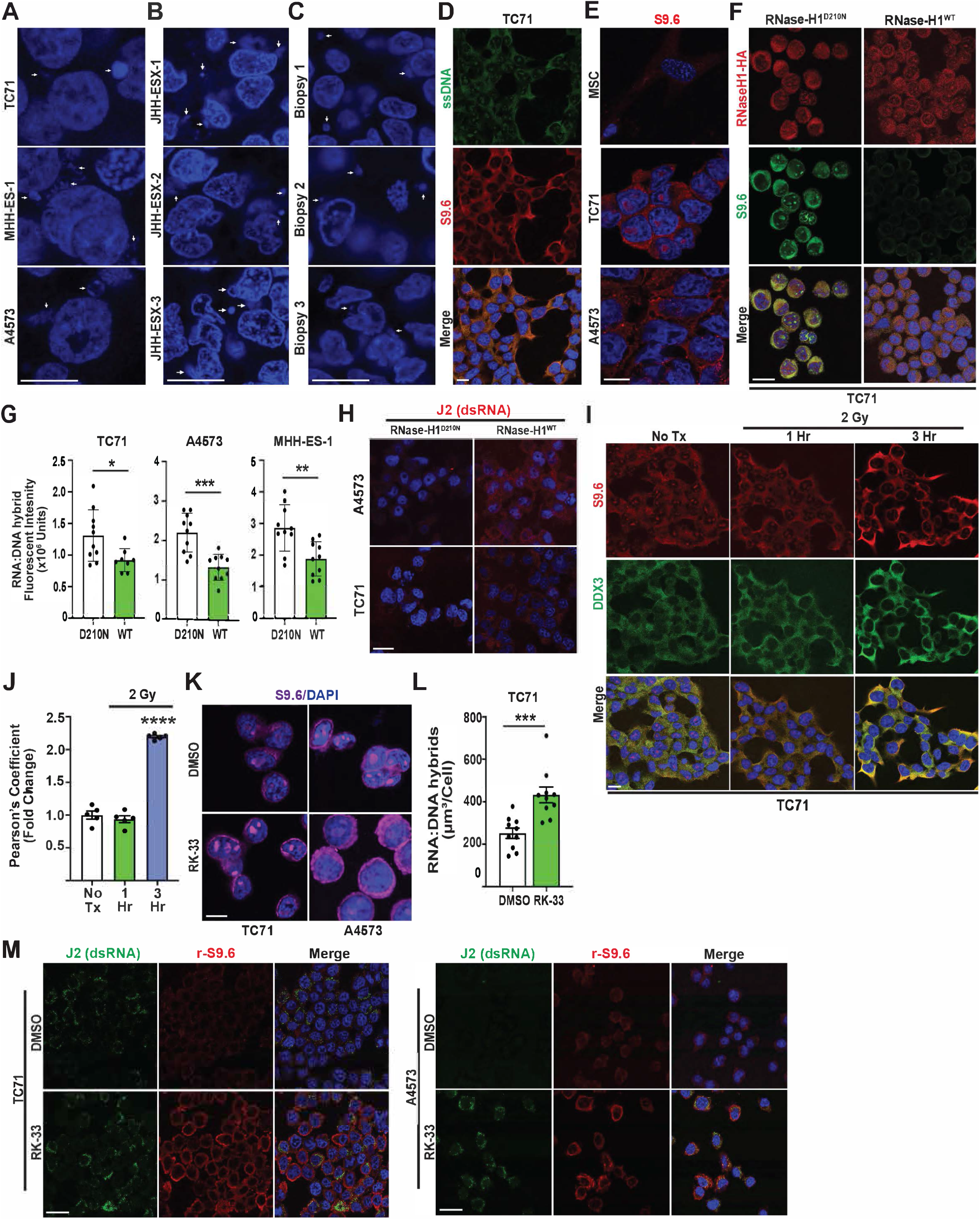
DDX3 interacts with and modulates cytoplasmic oligonucleotide substrates in Ewing sarcoma. (A, B, C) DAPI staining of dsDNA in 3 independent Ewing sarcoma cell lines (A), 3independent patient-derived xenografts (B), and 3 independent patient samples (C). White arrows show examples of extra-nuclear dsDNA substrates. Mag bars = 10 µm. (D) Immunofluorescent images of TC71 EWS cells. Data are representative of two independent experiments. Green = singlestranded DNA (ssDNA); Red = RNA:DNA hybrid structures; Blue = DAPI stain. Mag bar = 24 µm. (E) Representative images of S9.6 staining of umbilical chord-derived human mesenchymal stem cells (MSC), TC71, and A4573 EWS cell lines. Mag bar = 10 µm. (F, G) Three independent EWS cell lines, TC71, A4573, and MHH-ES-1, were transduced with lentivirus overexpressing either RNaseH1^WT^ or enzymatically-dead RNaseH1^D210N^ for 48 hours. Cells were then stained and immunofluorescent (IF) confocal images were obtained (F) and basal RNA:DNA hybrid abundance quantified (G). Results are representative of four to five hpf per condition from two independent experiments per cell line. Data represent mean fluorescent intensity per cell per hpf +/- SEM. *p<0.05, **p<0.01 and ***p<0.001 determined by unpaired t-tests. Red = HA-tag; Green = RNA:DNA hybrids (S9.6 staining); Blue = DAPI stain. Mag bar = 10 µm. (H) TC71 and A4573 EWS cell lines were transduced with either RNaseH1^WT^ or enzymatically-dead RNaseH1^D210N^ for 48 hours. Cells were then stained with J2 to visualize dsRNA. Representative IF confocal images are shown. Mag bar = 10 µm. (I, J) Representative IF images of TC71 cells at 1 and 3 hours post-2 Gy irradiation (I). Colocalization of DDX3 and RNA:DNA hybrid fluorescence was analyzed and quantified using Pearson’s coefficient. Data are representative of five high power fields (hpf) per cohort from one of three independent experiments (J). Data are mean SEM. ****p<0.0001 determined by One-Way ANOVA followed by Dunnett’s multiple comparisons test. Green = DDX3; Red = RNA:DNA hybrid structures; Blue = DAPI stain. Mag bar = 24 µm. (K, L) TC71 and A4573 cells were treated with either vehicle control (DMSO) or 2 µM RK-33. Immunofluorescent Z-stack confocal images were obtained (K). TC71 images were measured for volume of RNA:DNA hybrid structures per cell (L). Results are representative of five hpf per condition from two independent experiments. Data represent mean fluorescent intensity per cell +/- SEM. ***p<0.001 determined by unpaired t-test. Mag bar = 10 µm. (M) Representative doublestained IF images analyzing RNA:DNA hybrid and double-stranded RNA (dsRNA) distribution with a rabbit-S9.6 and J2 antibodies, respectively, in TC71 and A4573 EWS cell lines treated with either DMSO or 2 µM RK-33. Mag bar = 10 µm.

To query whether the interaction of DDX3 with cytoplasmic RNA:DNA hybrids could contribute to the radiosensitization effect of RK-33 in EWS, we treated EWS cell lines with 2 Gy IR and examined changes in the interactions between DDX3 and RNA:DNA hybrid substrates. DDX3 co localization with cytoplasmic RNA:DNA hybrid structures significantly increased in a time-dependent manner following IR (p<0.0001; Figure 5I, 5J), suggesting that IR damage increases the interaction of DDX3 with cytoplasmic RNA:DNA hybrids.

### Inhibition of DDX3 helicase activity by RK-33 impairs resolution of cytoplasmic RNA:DNA hybrid structures

DDX3 is an ATP-dependent DEAD-box RNA helicase that resolves oligonucleotide duplexes in a non-processive manner ^6^. The DDX3 inhibitor, RK-33, specifically binds to the catalytic pocket of the DDX3 ATPase catalytic domain, thus inhibiting helicase activity and preventing the resolution of oligonucleotide duplexes ^11, 12^. Thus, we hypothesized that impairment of DDX3 RNA helicase activity by RK-33 should result in an increase of hybrid structures. Indeed, when EWS cell lines were treated with RK-33 alone, S9.6 staining of cytoplasmic RNA:DNA hybrid structures significantly increased (p=0.0007; Figure 5K, 5L). We confirmed that the increased cytoplasmic staining with S9.6 reflects an increase in RNA:DNA hybrids and not an increase in dsRNA by co-staining EWS cell lines with the dsRNA-specific murine J2 antibody and the rabbit-derived S9.6 antibody (Figure 5M). The rabbit S9.6 antibody demonstrated a similar increase in cytoplasmic staining pattern in both TC71 and A4573 cell lines, as demonstrated using the murine hybridoma S9.6 antibody (Figure 5K), in response to RK-33 treatment (Figure 5M). Despite increased dsRNA staining following DDX3 inhibition, the J2 staining showed little to no co-localization with S9.6 moieties. Collectively, these data confirm that cytoplasmic RNA:DNA hybrids are present in EWS and demonstrate that inhibition of DDX3 RNA helicase activity attenuates the resolution of cytoplasmic RNA:DNA hybrid structures, thus increasing their abundance.

### DDX3 inhibition by RK-33 sequesters RAD51 in the cytoplasm following IR

Our data support the hypothesis that RNA:DNA hybrid structures provide a cytoplasmic scaffold for DDX3 in EWS, and that impairing the helicase activity of DDX3 with RK-33 further enhances this cytoplasmic localization. We next tested whether the cytoplasmic localization of DDX3 contributes to the radiosensitizing effect of RK-33. As described above, the HR proteins RAD51, RPA32, RECQL1, and XRCC2, which co-immunoprecipitate with DDX3 (Figure 4C), were also observed in the cytoplasmic fraction of two independent EWS cell lines (Figure 4D, Supplemental Figure 2B). The importance of subcellular localization and its effect on DDR of only one of these four candidate DDR proteins, RAD51, has been well characterized ^32, 48–51^. Accurate repair of damaged DNA depends on the proper localization of RAD51 to sites of dsDNA breaks following IR-induced damage ^52^. Additionally, RAD51-depletion impairs RAD51 loading in both homologous DDR and in the resolution of stalled replication forks ^52^. In both cases, single stranded regions of DNA are then exposed to exonucleases which ultimately result in increased genomic instability and accumulation of cytoplasmic self-DNA ^52, 53^. Considering that EWS has high basal levels of R-loops, replication stress ^18^ and cytoplasmic oligonucleotide structures (Figure 5A-C), we examined whether DDX3 helicase impairment also altered RAD51 cellular localization, providing a mechanistic explanation for our observation that RK-33 acts as a radiosensitizing agent.

Under basal conditions, RAD51 localizes to both the cytoplasm and nucleus and partially colocalizes with DDX3 in the cytoplasm in TC71 and A4573 EWS cells (Figure 6A). Within four hours of IR (2 Gy), distinct RAD51 nuclear foci were observed which localized with nuclear γ-H2A.X foci, whereas DDX3 did not (Figure 6B). In contrast, when DDX3 helicase activity was inhibited by RK-33, IR-induced RAD51 focus formation was diminished, and pronounced RAD51 colocalization with cytoplasmic DDX3 and RNA:DNA hybrids was observed (Figure 6C). Growing evidence suggests that RNA:DNA hybrids form at sites of DSBs and play a role in DDR ^54–57^. Additionally, the presence of RNA:DNA hybrids has been associated with recruitment of RAD51 to sites of DSBs ^55–57^. Therefore, we explored whether the increase of cytoplasmic RNA:DNA hybrids following IR and DDX3 helicase inhibition may be sequestering RAD51 in the cytoplasm. To test this, we overexpressed RNaseH1^WT^ in three independent EWS cell lines to resolve cytoplasmic RNA:DNA hybrids, and then quantified nuclear RAD51 foci following RK-33 treatment. If RAD51 is sequestered in the cytoplasm, we hypothesized that the percentage of total cellular RAD51 foci that is nuclear should decrease. In cells expressing the RNaseH1^D210N^ enzymatically inactive control, RAD51 foci increased with IR alone (Figure 6D) as expected from our data demonstrating IR-induced nuclear and cytoplasmic RAD51 foci formation (Figure 6B, C). Moreover, when RNaseH1^D210N^-expressing EWS cells were treated with both IR and RK-33, which increases cytoplasmic RNA:DNA hybrid structures, there was a significant <u>decrease</u> in nuclear RAD51 foci compared to cells treated only with IR in three independent EWS cell lines (Figure 6D – red vs blue bars). Importantly, when cytoplasmic RNA:DNA hybrids were resolved by overexpression of RNaseH1^WT^ (Figure 5F), RK-33 treatment combined with IR was unable to significantly alter the nuclear formation of RAD51 foci (Figure 6D – red vs blue bars), suggesting that removal of the cytoplasmic RNA:DNA hybrid scaffolds allowed nuclear localization of RAD51. These findings support our hypothesis that impairment of DDX3 helicase function, via RK-33, prevents resolution of DSBs induced by IR by sequestering RAD51 in the cytoplasm and preventing formation of nuclear foci at the site of DSBs through a mechanism that is dependent on modulation of cytoplasmic RNA:DNA hybrid structures.

**Figure 6.**
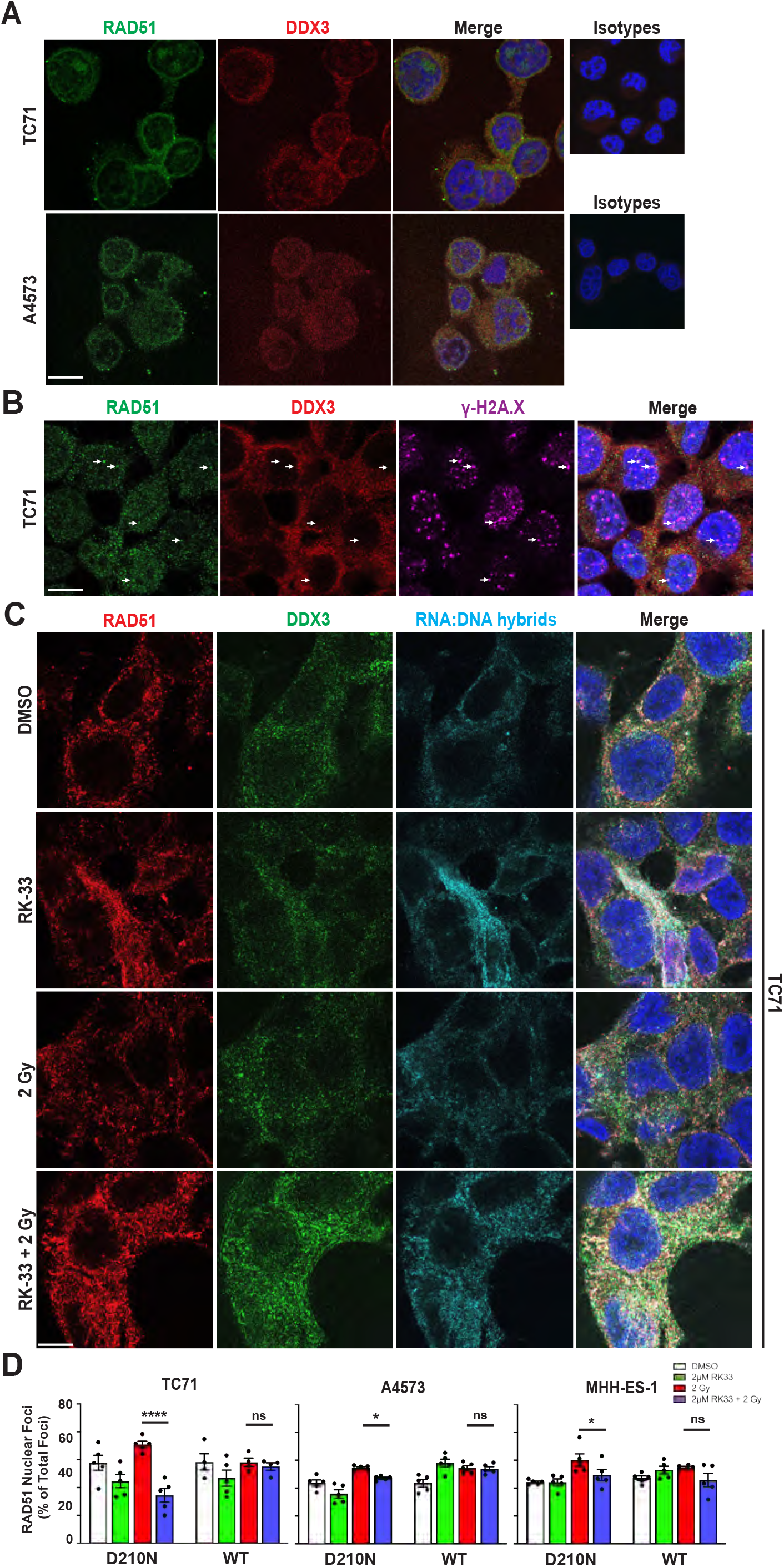
Inhibition of DDX3 helicase activity sequesters RAD51 in the cytoplasm in an RNA:DNA hybrid dependent manner following irradiation. (A) Representative immunofluorescent images of two EWS cell lines, TC71 and A4573, demonstrating cytoplasmic colocalization of endogenous RAD51 (green) and DDX3 (red) at basal levels. Data are representative of three independent experiments. Blue = DAPI stain. Mag bar = 10 µm. (B) Representative immunofluorescent images of TC71 cells demonstrating colocalization of RAD51 (green) with DSBs, evidenced by γ-H2A.X (purple) staining, 3 hours following 2 Gy RT. Data are representative of two independent experiments. Red = DDX3; Blue = DAPI stain. Mag bar = 10 µm. (C) Immunofluorescent images of TC71 cells at 3 hours following treatment with either DMSO (top), 2 µm RK-33 (second row), DMSO + 2 Gy irradiation (third row), or 2 µM RK-33 + 2 Gy (bottom). Data are representative of three independent experiments. Green = DDX3; Red = RAD51; Cyan = RNA:DNA hybrids; Blue = DAPI stain. Mag bar = 20 µm. (D) Three independent EWS cell lines, TC71, A4573, and MHH-ES-1, were transduced with lentivirus overexpressing either RNaseH1^WT^ or enzymatically-dead RNaseH1^D210N^ for 48 hours prior to performing radiosensitization assays. Cells were stained and immunofluorescent, Z-stacked confocal images were obtained. RAD51 foci were quantified for each experimental cohort. Results represent nuclear RAD51 foci as a percentage of total RAD51 foci from all Z-planes of 4-5 representative hpf from each experimental cohort of each cell line. Data represent one of three independent experiments. Data are mean ± SEM. *p<0.05 and ***p<0.001 determined by Two-Way ANOVA followed by Šídák’s multiple comparisons test.

## Discussion

In this study, we elucidate a novel mechanism of radiosensitization for the therapeutic targeting of cancers that accumulate cytoplasmic nucleic acid structures which arise in tumors with high levels of replication stress, like EWS. To our knowledge, we are the first to demonstrate that altering cytoplasmic RNA:DNA hybrid abundance via inhibition of a DEAD-box RNA helicase abrogates the repair of IR-induced DSBs by cytoplasmic sequestration of an essential DDR protein, thereby impeding DNA repair following radiotherapy (Figure 7). Inhibition of DDX3 helicase activity, using RK 33, increased cytoplasmic RNA:DNA hybrid structures. When RK-33 was combined with IR, RAD51 was sequestered in the cytoplasm and colocalized with cytoplasmic RNA:DNA hybrids, thereby reducing the formation of nuclear RAD51 foci required for HR of DSBs. Whether the increased cytoplasmic RAD51 abundance occurred from RAD51 binding to excised nuclear RNA:DNA hybrid structures before exportation out of the nucleus or from *de novo* binding to cytoplasmic RNA:DNA scaffolds, the sequestration phenotype was reversed following removal of RNA:DNA hybrid structures, thus demonstrating a role for these structures in DDR. Our data suggest that leveraging the by-products of increased replication stress, i.e. cytoplasmic RNA:DNA hybrid structures, as scaffolds for DDR protein sequestration could provide a novel and viable therapeutic approach for targeting cancers characterized by high levels of replication stress.

**Figure 7.**
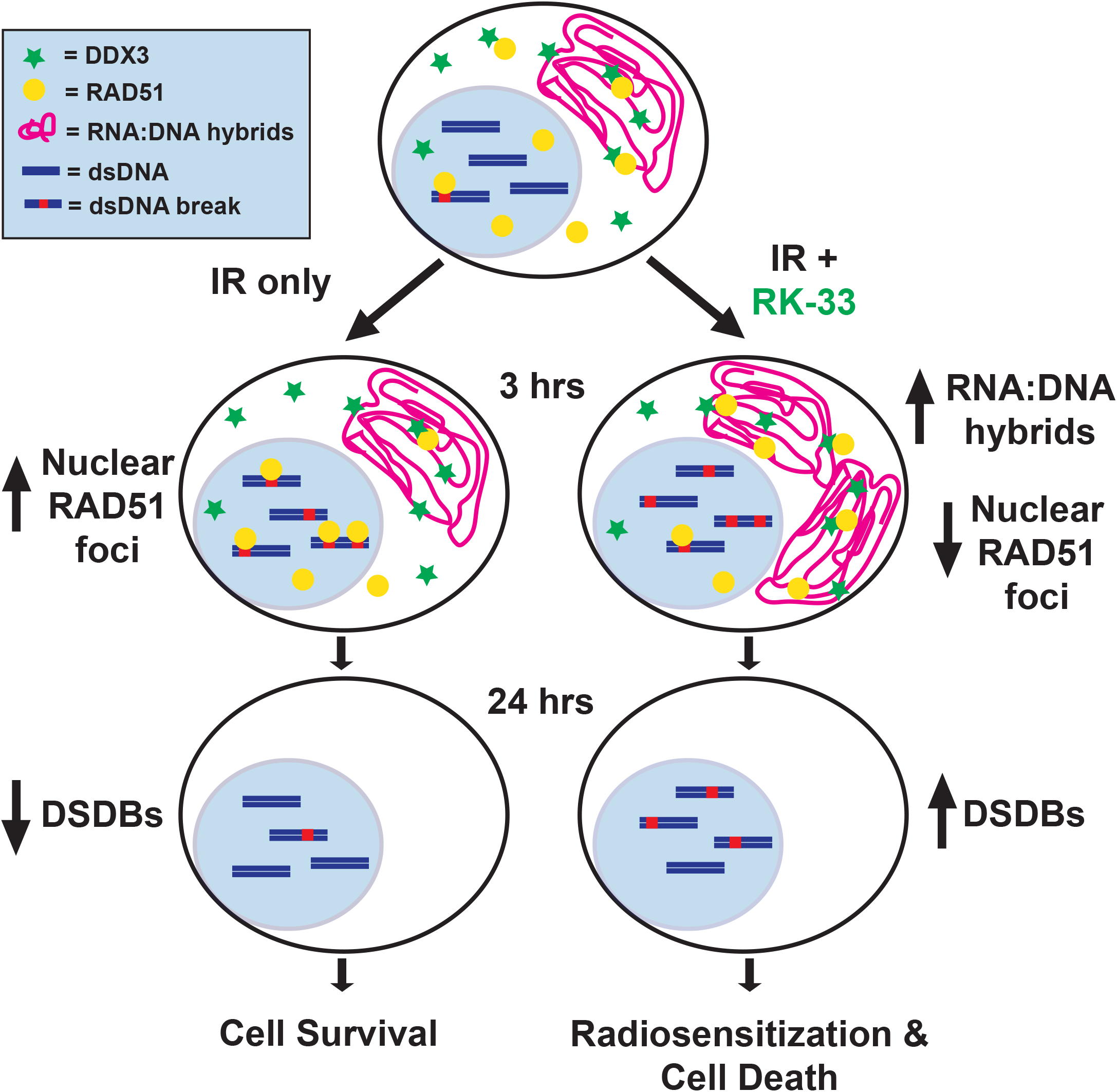
Mechanistic model of Ewing sarcoma radisosensatization following DDX3 RNA helicase inhibition. Ewing sarcoma’s genomic instability is evidenced by basal levels of double-stranded DNA breaks (DSBs) and cytoplasmic RNA:DNA hybrid structures. Endogenous DDX3 associates with both cytoplasmic RNA:DNA hybrids and RAD51. When ES is subjected to radiation, RAD51 foci form at sites of radiation-induced DSBs within 3 hours. Within 24 hours, the majority of DSBs are resolved back to basal levels resulting in cell survival. In contrast, inhibition of DDX3 RNA helicase activity, with RK-33, impairs resolution and increases the abundance of cytoplasmic RNA:DNA hybrid structures. RAD51 is sequestered in the cytoplasm in an RNA:DNA hybrid-dependent manner and results in decreased nuclear foci formation, thereby impairing homology directed repair of radiationinduced DSBs. The resultant radiosensitization significantly impairs cell survival over radiation therapy alone.

Accumulation of cytoplasmic self-DNA structures is one of the many cellular sequelae of neoplastic hyperproliferation, *i.e.* oncogene-induced replication stress [reviewed^58^], which is characteristic of EWS biology ^18^. We confirmed that not only cytoplasmic DNA substrates, but also ssDNA and RNA:DNA hybrid structures, are present in EWS cell lines and PDX models, as well as in primary human EWS samples. Indeed, ssDNA and RNA:DNA hybrid structures were present in both the nucleus and cytoplasm. We demonstrated a robust association between DDX3 and the cytoplasmic RNA:DNA hybrid structures compared to ssDNA substrates. Several studies have implicated DDX3 in cellular pathways which regulate cytoplasmic oligonucleotide moieties. DDX3 has been shown to act as a sensor of cytoplasmic ssRNA and dsRNA through the RIG-1 (retinoic acid inducible gene 1)/ MAVS (mitochondrial antiviral signaling protein) signaling pathway which activates IKKε (IκB kinase-ε)/IRF3 (interferon regulatory factor 3) signaling to induce a type 1 interferon response ^59–62^. Roles for DDX3 have also been described in the innate immunity pathways involving the NF-κB (nuclear factor-κB) pathway through IKKβ ^63^; the Toll-like receptor 7/8 signaling through NIK (NF-κB-inducing kinase)/ IKKα and alternative NF-κB signaling pathways ^64^. Additionally, a new study suggests that impairing the helicase activity of DDX3 results in a cytoplasmic accumulation of dsRNA, activating a dsRNA-sensing pathway through MDA5 (melanoma differentiation-associated gene 5) signaling to induce an intrinsic type 1 interferon response in breast cancer ^65^. Our findings expand the biological spectrum of cytoplasmic DDX3 function to include regulation of RNA:DNA hybrid structures. Importantly, our data indicate that DDX3 RNA helicase activity modulates and resolves cytoplasmic RNA:DNA hybrid structures *in vivo* in tumors with high levels of replication stress. In the context of IR, both RNA:DNA hybrid abundance and DDX3 localization to cytoplasmic hybrid structures increased in a time-dependent manner, while RK-33 impairment of DDX3 helicase activity further amplified the abundance of cytoplasmic RNA:DNA hybrids. This increase in RNA:DNA hybrid structures provided a cytoplasmic scaffold for sequestering the DDR protein RAD51.

These findings are based on immunofluorescent analysis of cytoplasmic oligonucleotides using antibody S9.6, which detects RNA:DNA hybrids, and J2, which binds dsRNA. There are reports in the literature suggesting S9.6 may not be specific for RNA:DNA hybrids and may, in fact, also recognize dsRNA in formalin fixed cells. In our cell lines and under our experimental conditions, it appears that S9.6 and J2 detect different substrates, with minimal cross-reactivity of S9.6. Evidence for this includes a) the ability of wild-type, but not catalytically inactive, RNase-H1 to eliminate the S9.6 substrate (Figure 5F), b) the inability of wild-type RNase-H1 to eliminate the J2 substrate (Figure 5H), and c) the lack of overlap of the murine J2 and rabbit S9.6 immunofluorescence signals in co immunofluorescence experiments (Figure 5M). This constellation of results is most consistent with our contention that EWS cell lines accumulate cytoplasmic RNA:DNA hybrids, and support our model that DDX3 regulates the abundance of cytoplasmic RNA:DNA hybrids, regulating the subcellular localization of RAD51.

Mounting evidence supports the importance of cellular localization of DDR proteins in preserving or impairing genomic stability ^32–37, 48–51, 66, 67^. Indeed, cytoplasmic sequestration of DDR proteins is an emerging therapeutic approach for cancer patients. Of note, previous studies revealed that lapatinib, an epidermal growth factor receptor (EGFR) inhibitor, induced a cytoplasmic localization of both BRCA1 and EGFR in triple-negative breast cancer models. This cytoplasmic sequestration resulted in a synthetic lethality when combined with a PARP inhibitor ^67^. Leveraging this cytoplasmic sequestration phenomenon, a Phase 1 clinical trial was designed for patients with metastatic triple-negative breast cancer (NCT02158507) which demonstrated ‘promising antitumor activity’ in the enrolled patients ^68^. Other groups have implicated cytoplasmic sequestration of RAD51 in contributing to genomic instability and impaired DNA-damage repair through mechanisms involving AKT1 signaling ^32^, IGF-R1 signaling ^50^, and hexavalent chromium ^69^. We discovered a novel mechanism for the cytoplasmic sequestration of DDR proteins where DDX3 helicase inhibition sequesters RAD51 predominantly in the cytoplasm, reduces the formation of nuclear RAD51 foci, and thus prevents RAD51 from participating in the nuclear repair of dsDNA breaks following IR. Our findings align with a recent study that also demonstrated impairment of nuclear RAD51 focus formation in sarcomas with high levels of HR deficiency ^70^. Importantly, when we tested this therapeutic approach *in vivo* (Figure 3), pronounced tumor ablation resulted which was maintained for several weeks. Therefore, our pre-clinical data suggest that physically altering the cellular localization of DDR proteins, such as RAD51, is a novel way to potentiate the effectiveness of radiation therapy, a critical treatment modality for many cancer patients.

The RNA helicase DDX3, which our data show plays a pivotal role in regulating the subcellular localization of RAD51, has been studied in multiple adult and pediatric cancers ^7, 12, 23, 71–74^. The role of DDX3 in these cancers is context-dependent, acting as either an oncogene or a tumor suppressor depending on the cancer and even cancer sub-types [reviewed by ^23^]. Previously, we have demonstrated that DDX3 functions as an oncogene in EWS ^10^. Here we demonstrate that 75% of EWS tumors examined express moderate to high levels of DDX3, and that high levels of DDX3 expression are associated with poor survival. These findings, in combination with our previous work, strongly implicate the RNA helicase DDX3 as a therapeutic target for EWS patients.

EWS patients with recurrent, metastatic and unresectable disease currently have poor prognoses for long-term survival ^1, 2^. Considering that radiation therapy is commonly used to treat these patients ^14, 15^, our finding that the DDX3 RNA helicase inhibitor, RK-33, functions as a radiosensitizing agent is particularly important. We found that DDX3 inhibition, both genetically and chemically, sensitizes EWS to IR-induced cell death by delaying the repair of IR-induced DSBs in multiple EWS cell lines. Clonogenic survival of EWS cells was impaired compared to IR or RK-33 alone *in vitro* when treated with 2 µm RK-33 and 2 Gy IR. In agreement with these findings, a pronounced *in vivo* tumor ablation occurred following RK-33 radiosensitization in a DDX3^high^ EWS PDX model, thus demonstrating the clinical potential for RK-33 as a radiosensitizing agent for EWS patients.

Although the complete array of mechanisms by which DDX3 enhances radiotherapy efficacy remains largely unknown, our findings suggest that modulation of the subcellular localization of RAD51 is a key part of this process. Impairment of DDX3 by RK-33 has previously been shown to radiosensitize prostate ^71^, medulloblastoma ^75^ and lung cancer ^12^, but the mechanisms behind these observations have not been elucidated. One study found that RK-33 radiosensitization in breast cancer involves abrogation of mitochondrial translation with concurrent increases in reactive oxygen species ^24^. In the context of human hepatocellular carcinoma, *in vivo* genomic loss of DDX3X in liver tissue resulted in increased single-strand break and DSB signaling, as well as decreased expression of nucleotide excision repair proteins that resulted in liver tumorigenesis ^76^. These studies indicate an important role for DDX3 in maintaining genomic stability through mechanisms of DDR. Recent studies have also implicated a role for DDX3 function in NHEJ repair of DSBs ^12^. When we examined DDR pathways affected by DDX3 function, we found that both NHEJ and HR repair were impaired with siDDX3 knockdown. In contrast to previous studies where translation and protein expression were altered following DDX3 impairment, we detected no gross alterations in HR/NHEJ DDR protein abundance nor double-strand break signaling following RK-33 treatment in EWS. Instead, co immunoprecipitation experiments revealed protein-protein interactions between DDX3 and select HR proteins but not proteins involved in NHEJ. Contrary to what we expected, endogenous DDX3 did not colocalize with DSBs but rather demonstrated a predominant cytoplasmic localization, suggesting that DDX3 does not robustly interact with DDR proteins at the site of DNA repair and is thus unlikely to contribute to DDR at sites of DSBs.

In conclusion, we report a novel mechanism of radiosensitization whereby DDR is impaired due to cytoplasmic sequestration of RAD51 following the inhibition of DDX3 RNA helicase activity. Cytoplasmic sequestration of RAD51 was dependent on the presence of cytoplasmic RNA:DNA hybrid structures. Importantly, we found that DDX3 helicase activity plays an active role in modulating cytoplasmic RNA:DNA hybrid levels, thus indirectly regulating RAD51 subcellular localization, and thereby inducing radiosensitization of EWS. These data further emphasize the importance of DDR protein localization in cancer biology and the need to develop new therapeutics directed at manipulating DDR protein localization. Additional studies are needed to determine whether a cytoplasmic DDX3-RNA:DNA hybrid-RAD51 complex is also present in other solid tumors which can potentially be targeted with radiation therapy and RK-33 treatment. Indeed, therapies aimed at leveraging the biological effects of genomic instability hold promise for increasing the potency and efficacy of clinical radiation therapy in multiple cancer types.

## Acknowledgments

We thank N, Ghazale, J.Biswas, R.Singer, and T. Bowman for the generous donation of the ubc-Hu-RNASEH1-3xHA and ubc-Hu-RNASEH1-D210N-Halo-3xHA lentiviral vectors; T. Bowman and A. Baker for helpful insights and discussions; H. Guzik and V. DesMarais of AECOM Analytical Imaging Facility for training and technical assistance, supported by NCI cancer center support grant P30CA013330. This work was funded in part by Alex’s Lemonade Stand Foundation and by the Montefiore Einstein Cancer Center Support Grant Number 2P30CA013330. V.R. was supported by a grant from FAMRI. A.J.R.B. was supported by the National Institutes of Health R01CA152063 and 1R01CA241554; the Cancer Prevention and Research Institute of Texas RP150445; and the Stand Up 2 Cancer-Cancer Research UK RT6187. A.G. was supported by American Association for Cancer Research-AstraZeneca Stimulating Therapeutic Advances through Research Training grant 18-40-12-GORT. ABTR14B2-Q was supported by NCTN Operations Center Grant U10 CA180886, Human Specimen Banking Grant U24 CA114766, and NCTN Statistics & Data Center Grant U10 CA 180899 of the Children’s Oncology Group from the National Cancer Institute of the National Institutes of Health. Additional support for research was provided by a grant from the WWWW (QuadW) Foundation, Inc. (www.QuadW.org) to the Children’s Oncology Group. Disclaimer: The content is solely the responsibility of the authors and does not necessarily represent the official views of the National Institutes of Health.

## Author Contributions

Conceptualization, M.E.R., M.A., B.A.W., V.R., A.J.R.B., and D.M.L.; Methodology, M.E.R., M.A., A.G., B.A.W., V.R., A.J.R.B., and D.M.L.; Validation, M.E.R., M.A., A.G., R.W., B.A.W., J.W., and N.tH.; Formal Analysis, M.E.R., M.A., A.G., and B.A.W.; Investigation, M.E.R., M.A., A.G., R.W., B.A.W., J.W., P.C., and N.tH.; Resources, P.J.vD., V.R., A.J.R.B., and D.M.L.; Writing – Original Draft, M.E.R., M.A., A.G., and D.M.L.; Writing –Review & Editing, M.E.R., M.A., A.G., B.A.W., P.J.vD., V.R., A.J.R.B., and D.M.L; Visualization, M.E.R., M.A., A.G. and R.W.; Supervision, P.J.vD., A.J.R.B., and D.M.L.; Funding Acquisition, A.G., P.J.vD., V.R., A.J.R.B., and D.M.L..

## STAR Methods

### Cell lines and Xenografts

Established EWS cell lines TC71 (RRID:CVCL_2213) and TC32 (RRID:CVCL_7151) were acquired from Children’s Cancer Respository (https://cccells.org), A4573 (RRID:CVCL_6245) was a kind gift from the laboratory of Katia Scotlandi, and MHH-ES-1 (ACC-167; RRID:CVCL_1411) was purchased from German Collection of Microorganisms and Cell Cultures (dsmz.de, Braunschweig, Germany). Cell lines were validated using short-tandem repeat (STR) profiling at Albert Einstein College of Medicine’s (AECOM) Department of Genetics Genomic Core. HEK293T cells (CRL-157; RRID:CVCL_0063) were purchased from American Type Culture Collection (ATCC, Manassas, VA, USA). U2OS cells containing an integrated copy of each reporter and the homing endonuclease I*Sce*I in pCAGGS vectors with control were kind gifts from Dr. Maria Jasin (Memorial Sloan Kettering Cancer Center, New York, USA) and Dr. Jeremy Stark (City of Hope, California, USA). DDX3 shRNA knockdown cell lines were generated as previously described by Wilky *et al.* (2016). EWS-4 PDX was a gift from Chand Khanna (National Cancer Institute, Bethesda, MD) while the JHH-ESX-1, JHHESX-2, and JHH-ESX-3 xenografts were generated in our laboratory on a tumor banking protocol approved by Johns Hopkins University School of Medicine IRB. EWS xenografts were coated with Matrigel basement membrane (BD Biosciences, Bedford, MA, USA) and implanted subcutaneously into NOD.Cg-*Prkdc*^scid^ *Il2rg*^tm1Wjl^/SzJ (NSG, Strain #:005557; RRID:IMSR_JAX:005557) mice purchased from The Jackson Laboratory (Bar Harbor, ME, USA). All PDXs were passaged subcutaneously in NSG mice at least 3 times prior to experimental usage.

### Cell Culture

Cell lines were cultured in RPMI 1640 medium (Gibco-ThermoFisher Scientific, Waltham, MA, USA) containing 10% heat-inactivated fetal bovine serum (Atlanta Biologicals, Flowery Branch, GA, USA) at 37°C in 5% CO^2^. Cell lines were passaged between 6 to 10 times using 0.25% trypsin-EDTA (Gibco-ThermoFisher Scientific) between thawing and experimental collection. ATTC Universal Mycoplasma Detection Kit (ATCC 301012K, Fischer Scientific, Waltham, MA) was used per manufacturer’s instructions to screen cultures every 3 to 6 months to confirm absence of mycoplasma infection.

### Radiosensitization Assay

EWS cells were either grown in 4 well chamber slides (Nunc Lab-Tek II Chamber Slide^™^ system, Thermo Scientific, Hudson, NH, USA) at a density of 140,000 cells/well or in 6 well plates at densities ranging from 2.5 to 5 x 10^5^ cells on coverslips coated with bovine collagen type I (Gibco-Thermo Fisher Scientific) and incubated for 24 hours at 37°C and 5% CO^2^. RK-33 (CAS# 1070773-09-9) was dissolved in dimethyl sulfoxide (DMSO, Sigma-Aldrich) and added to the cultured cells 45-60 minutes prior to IR (2 Gy). The cells were maintained at 37^°^C and 5% CO^2^ until the desired timepoint.

### Clonogenic Assays

Clonogenic assays were performed as previously described ^77^. Briefly, radiosensitization assays were performed using EWS cells as described above. Six hours following IR, cells were detached from the plate using 0.25% trypsin-EDTA (Gibco-ThermoFisher Scientific). Clonal densities of 400 or 800 cells/well in 6 well plates containing conditioned media were then plated in triplicate. Cells were then grown at 37°C and 5% CO^2^ for 5 days undisturbed. Clones were then fixed with 4% paraformaldehyde (Thermo Scientific) at room temperature for 30 minutes and stained with 0.5% crystal violet in methanol. Clones were imaged using the ChemiDoc Touch Imaging System (RRID:SCR_021693, Bio-Rad Laboratories) and numbers of clones were quantified using ImageJ2, version 2.3.0/1.53f software (RRID:SCR_003070, NIH, Bethesda, MD). The analysis of clone numbers was performed blinded.

### Immunofluorescence

For immunofluorescent staining, cells were fixed with 4% paraformaldehyde (Thermo Scientific) for 10 mins, washed 1x with phosphate buffered saline (PBS), permeabilized using 0.2% Triton-X in PBS for 15 mins, then blocked for 30 minutes using 5% goat serum, 1% bovine serum albumin (BSA), 0.2% Triton-X PBS blocking buffer. Slides were then incubated with primary antibodies (Supplemental Table 1) diluted in blocking buffer overnight at 4°C. Slides were washed 3 times with 0.1% Tween™ 20 (Thermo Fisher Scientific) PBS the next day followed by 1 hour incubation at room temperature with fluorophore labeled secondary antibodies (Supplemental Table 1) diluted in blocking buffer. Slides were then washed 3x with PBS and mounted the coverslips to slides using ProLong™ Diamond Antifade Mountant with diamidino-2-phenylindole (DAPI) (Thermo Fisher Scientific). Slides were imaged using AECOM’s Analytical Imaging Facility’s Leica SP5 Acousto-Optical Beam Splitter (Leica Microcystems, Wetzlar, Germany) confocal microscope. Adobe Photoshop 2021 (RRID:SCR_014199, Adobe, San Jose, CA, USA) was used to globally process images for contrast, size, and brightness. Images were quantified using either ImageJ2 software (RRID:SCR_003070, NIH) or Volocity® software (RRID:SCR_002668, Quorum Technologies, Puslinch, Ontario, Canada).

### Tissue Microarray Immunohistochemistry

The EWS tissue microarrays were provided by the Children’s Oncology Group through project ABTR14B2-Q. Following deparaffination in xylene and the samples were rehydrated in decreasing ethanol dilutions. Endogenous peroxidase activity was blocked by endogenous peroxidase from Novolink Polymer Detection System (Leica Microsystems, Eindhoven, The Netherlands) and was followed by antigen retrieval by boiling for 20 minutes in EDTA buffer (pH 9.0). Slides were blocked with protein block from Novolink Polymer Detection System and subsequently incubated in a humidified chamber for 1 hour with anti-DDX3 (1:50, mAb AO196, RRID:AB_2936197, Sigma Aldrich)^78^. Post primary block, secondary antibodies and diaminobenzidine treatment were performed with the same Novolink Polymer Detection System according to the manufacturer’s instructions. The slides were lightly counterstained with hematoxylin and mounted. Images were scored for staining intensity using a scale of 0 to +3 by Dr. Paul J. van Diest, University Medical Center Utrecht, The Netherlands. The slides were scanned using a Hamamatsu, NanoZoomer XR C12000-21/-22.

### DNA Damage Repair Assays

The well-established DR-GFP, EJ5-GFP, and SSA-GFP reporter assays ^19, 20^ were used to evaluate the impact of DDX3X loss on homologous recombination, non-homologous end joining repair, and single-strand annealing repair, respectively, of induced double strand breaks. U2OS cells, containing an integrated copy of each reporter, were transfected with either scrambled control or DDX3X siRNA and seeded in 24-well plates using FuGENE® HD Transfection Reagent (Promega, Madison, WI, USA) as per manufacturer’s instructions. Twenty-four hours later, media was refreshed and cells were transfected with the I*Sce*I-pCAGGS vector or empty vector. Media was again refreshed after 12 hours and cells were allowed to grow for three more days. Samples were harvested and evaluated for percentage of GFP-positive cells using BD FACSCanto flow cytometer (BD Biosciences, Bedford, MA, USA). The experiment was conducted with triplicate independent transfections and analyzed using GraphPad Prism 9.3.1 (RRID:SCR_002798, San Diego, CA, USA).

### Western Blotting

Total cellular protein was extracted from cells and tumor tissue using the RIPA Lysis Buffer System (Santa Cruz Biotechnology, Dallas, TX, USA) as per manufacturer’s instructions. Protein concentration was determined utilizing the RC DC™ Protein Assay (Bio-Rad Laboratories, Hercules, California, USA) as per the manufacturer’s instructions. Serial dilutions of bovine serum albumin (BSA, Invitrogen-Thermo Fisher Scientific) were used to generate standard curves for protein quantification of experimental lysates. Samples were run on NuPAGE™ 4–12% Bis-Tris Protein gels (Invitrogen-Thermo Fisher Scientific) and transferred onto methanol-activated Immun-Blot® PVDF (polyvinylidene difluoride) Membranes (Bio-Rad Laboratories), followed by 1 hour blocking in 2% BSA in Tris-Buffered Saline, 0.1% Tween 20 Detergent (TBST, Sigma-Aldrich, St. Louis, MO, USA). Membranes were incubated with primary antibodies (Supplemental Table 1) in 2% BSA TBST blocking solution overnight at 4°C and subsequently incubated with HRP-conjugated secondary antibodies (Supplemental Table 1) diluted 1:10,000 in 4% BSA TBST for 1 hour at room temperature. Blots were developed using Clarity™ Western ECL Substrate (Bio-Rad Laboratories) and imaged using ChemiDoc Touch Imaging System (RRID:SCR_021693, Bio-Rad Laboratories). Antibodies were then stripped from blots using Restore Western Blot Stripping Buffer (Thermo Scientific) prior to incubation with additional primary antibodies. Adobe Photoshop 2021 (RRID:SCR_014199, Adobe) was used to globally process images for contrast, size, and brightness. Densitometry analysis was performed using ImageJ2 software (RRID:SCR_003070, NIH).

### Immunoprecipitations

To perform immunoprecipitation of whole cell lysates for DDX3, mouse anti-DDX3 IgG^2b^ (C-4, RRID:AB_10844621, Santa Cruz Biotechnology) or rabbit anti-DDX3 IgG (RRID:AB _2910140, Abcam, Waltham, MA, USA) were conjugated to SureBeads™ Protein G (1614023) or Protein A (1614013) Magnetic Beads (Bio-Rad Laboratories), respectively. Conjugation was performed per manufacturer’s instructions. Following conjugation of anti-DDX3 antibodies to the magnetic beads, whole cell lysates containing 500 µg of protein were applied to the beads and immunoprecipitation was performed using the manufacturer’s instructions at 4°C. Western blotting was performed in parallel where rabbit primary antibodies (Supplemental Table 1) were applied to blots containing samples from the mouse anti-DDX3 IgG^2b^ (C-4, RRID:AB_10844621, Santa Cruz Biotechnology) pulldown and mouse primary antibodies (Supplemental Table 1) were applied to blots containing the rabbit anti-DDX3 IgG (RRID:AB _2910140, Abcam, Waltham, MA, USA) pulldown samples.

### Subcellular Fractionations

Isolation and collection of cytoplasmic and nuclear cellular fractions of EWS cell lines was performed using NE PER Nuclear and Cytoplasmic Extraction Kit (PI78833, Thermo Scientific). Subcellular isolation of cytoplasmic and nuclear fractions was performed per manufacturer’s instructions. In parallel, whole cell lysates were generated, as described above, per experiment. Western blots were then performed, as described above, for experimental analysis of each fraction.

### Lentiviral Isolation and Transduction of RNaseH1 WT and RNaseH1 D210N

Lentiviral vectors ubc-Hu-RNASEH1-3xHA (RNaseH1 WT) and ubc-Hu-RNASEH1-D210NHalo-3xHA (RNaseH1 D210N) were gifts from Dr. Teresa Bowman. Lentiviral particles were generated by transfecting HEK293T with RNaseH1 WT or RNaseH1 D210N using FuGENE® HD Transfection Reagent (Promega) per manufacturer’s instructions. Viral supernatants were collected from transfected cultures and incubated in a 1:10 solution of 50% PEG 8000 and 1.5 M NaCl overnight at 4°C. Viral particles were then isolated via centrifugation, resuspended in serum-free RPMI 1640 media, and either used immediately or stored at -80°C. EWS cell lines were transduced with either RNaseH1^WT^ or RNaseH1^D210N^ lentiviral particles for 48 hours at 37°C in 5% CO^2^ prior to performing radiosensitization assays, described above.

### Animal studies

All mice procedures were approved by the Johns Hopkins Animal Care and Use Committee. Female, 3- to 6-month-old NOD.Cg-*Prkdc*^scid^ *Il2rg*^tm1Wjl^/SzJ (Strain #:005557; RRID:IMSR_JAX:005557) mice (JHU breeding colony, Baltimore, MD, USA) were used for experiments, with sample size of 10 randomly selected mice per cohort. Freshly isolated 3 mm^3^ EWS 4 and JHHES-X3 xenograft fragments coated with Matrigel (BD Biosciences) were implanted in the subcutaneous flanks. Once palpable, 7-9 mm^3^, animals bearing xenografts were randomly divided into 4 cohorts depending on treatments received: control (DMSO only), DMSO+IR, RK33 only, or RK33+IR). Either 50 mg/kg RK-33 or equivalent volume of 50 μl DMSO was injected intraperitoneal every other day for one week for a total of 3 treatments. After which, mice were exposed to 10 Gy IR 6 hours following the third injection using a small animal radiation research platform (SARPP) in the institutional Experimental Irradiators Core. Three more injections of DMSO or RK-33 every other day were administered in the following week post-IR. Tumor dimensions were measured twice weekly using calipers until reaching a diameter of 15 mm. Volumes were calculated using an elliptical formula and normalized to the initial tumor volume. No blinding was employed.

### Statistical Analysis

Data were analyzed for statistical significance using GraphPad Prism software: Version 9.3.1(RRID:SCR_002798, GraphPad Software, San Diego, CA). A confidence level 0.05 or less was considered statistically significant for all analyses. Kaplan-Meier curves were analyzed for significance using the log-rank (Mantel-Cox) test using R2 Genomics Platform (https://r2.amc.nl). Unpaired Student’s t test was used when comparing two independent data sets. To compare the results of multiple data sets across one variable, data were analyzed using one-way analysis of variance with either Dunnett’s or Šídák’s multiple comparisons posttests. Two-way analysis of variance with Šídák’s multiple comparisons posttest was applied to data sets with multiple comparisons across two variables. Repeated measures two-way analysis of variance with Tukey’s multiple comparisons posttest was utilized to analyze the *in vivo* PDX growth data at day 21.

### Data Availability

The data generated in this study are available upon request from the corresponding author.

## Supplemental Figures/Table Legends

**Supplemental Figure 1. RK-33 radiosensitization of EWS does not grossly alter expression of DDR proteins**

Western blot analysis of multiple DDR proteins from both the HR and NHEJ repair pathways. Cell lysates were collected from two independent EWS cell lines, MHH-ES-1 and TC71. Cells were treated with either DMSO, 1 µM RK-33, 2 Gy, or 1 µM RK-33 + 2 Gy and collected at either one- or twenty-four-hours post-treatment. Results are representative of 2-3 independent experiments per cell line.

**Supplemental Figure 2. DDX3 interacts with homologous repair proteins in multiple EWS cell lines**

(A) Immunoprecitipation (IP) of A4573, MHH-ES-1 and TC71 EWS cell lysates using antiDDX3 antibodies conjugated to magnetic beads. Western blots demonstrate co-immunoprecipitation of various DDR proteins with endogenous DDX3. Iso IgG = immunoprecipitation of TC71 cell lysates using control isotype antibodies. Results are representative of 3 independent experiments. (B) Subcellular fractionations of A4573 cells demonstrate the presence of DDX3, RAD51, RECQL1, RPA32 and XRCC2 in both cytoplasmic and nuclear compartments. Results are representative of 3 independent experiments.

**Supplemental Figure 3. DDX3 associates with cytoplasmic single-stranded DNA**

Immunofluorescent images of TC71 EWS cells at 1- and 3-hours post-2 Gy IR. Results are representative of two independent experiments. Green = single-stranded DNA (ssDNA); Purple = DDX3; Blue = DAPI stain. Mag bar = 24 µm.

**Supplemental Table 1. Reference lists of antibodies used in immunofluorescent staining and western blot experiments**

AF = AlexaFluor; ATM = ataxia telangiectasia mutated; ATR = ataxia telangiectasia and RAD3related protein; BRCA1 = breast cancer type 1 susceptibility protein; Chk1 = checkpoint kinase 1; DDX3 = DEAD box RNA helicase DDX3X; eIF4E = eukaryotic translation initiation factor 4E; H2A.X = histone H2A.X; HA = human influenza hemagglutinin; HRP = horseradish peroxidase; IF = Immunofluorescence; IgG = immunoglobulin G; Ku70 = ATP-dependent DNA helicase II 70 kDa subunit; Ku86 = ATP-dependent DNA helicase II 80 kDa subunit; Mre11 = meiotic recombination 11 homolog 1; NUP205 = Nuclear pore complex protein Nup205; p- = phosphorylated-; polyIgG = polyclonal IgG; RAD50 = DNA repair protein RAD50; RAD51 = DNA repair protein RAD51 homolog 1; RAD52 = DNA repair protein RAD52 homolog; ssDNA = single-stranded deoxyribonucleic acid; RPA32 = replication protein A subunit 32; WB = western blot; XPG = xeroderma pigmentosum group G-complementing protein; XRCC2 = X-ray repair cross-complementing protein 2; XRCC4 = X-ray repair cross-complementing protein 4.

