## Supplementary figures and images for "RNA Helicase DDX3 Regulates RAD51 Localization and DNA Damage Repair in Ewing Sarcoma"

### Supplemental Figure 1

#### Supplemental Figure 1

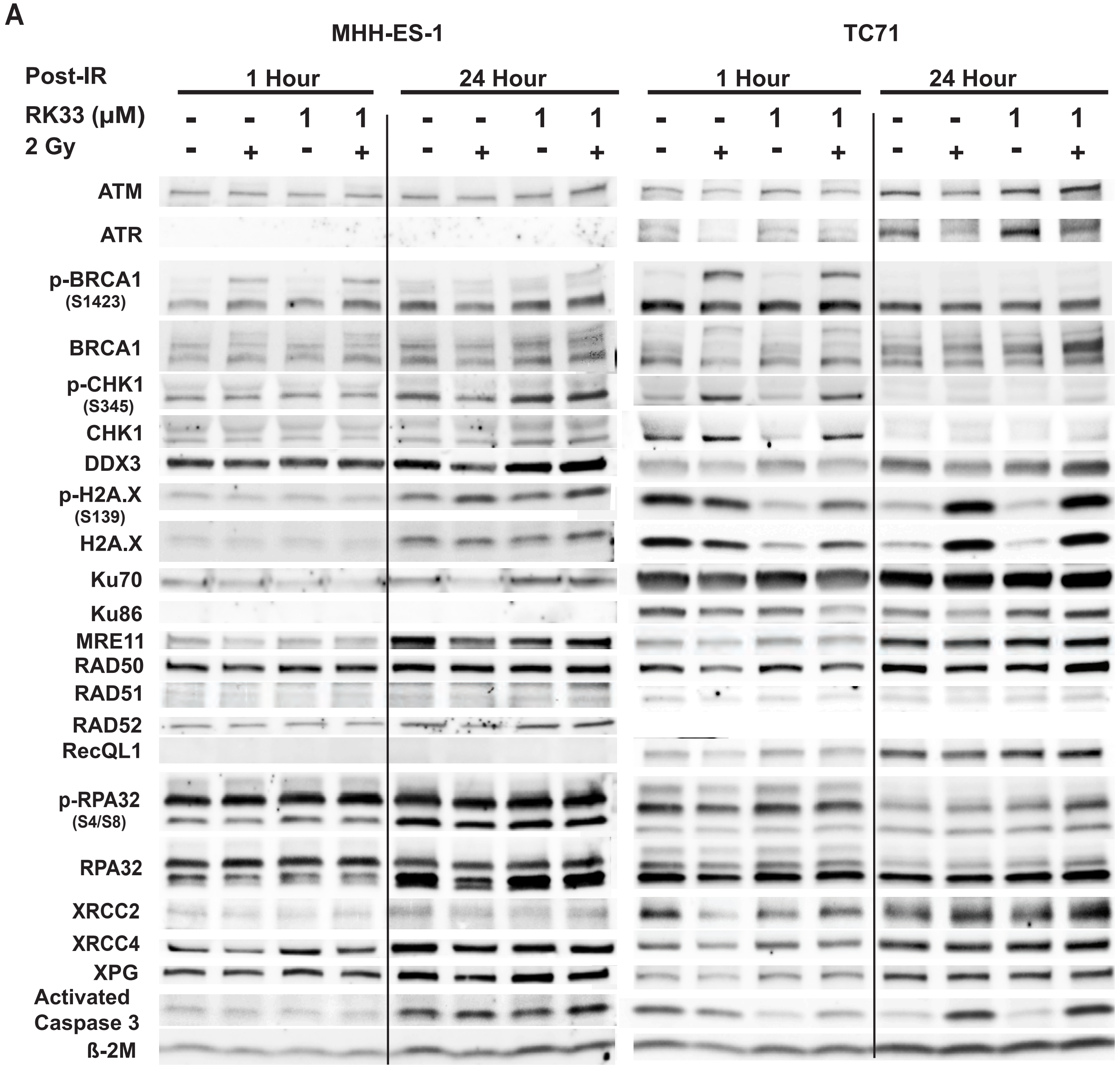

### Supplemental Figure 2

#### Supplemental Figure 2

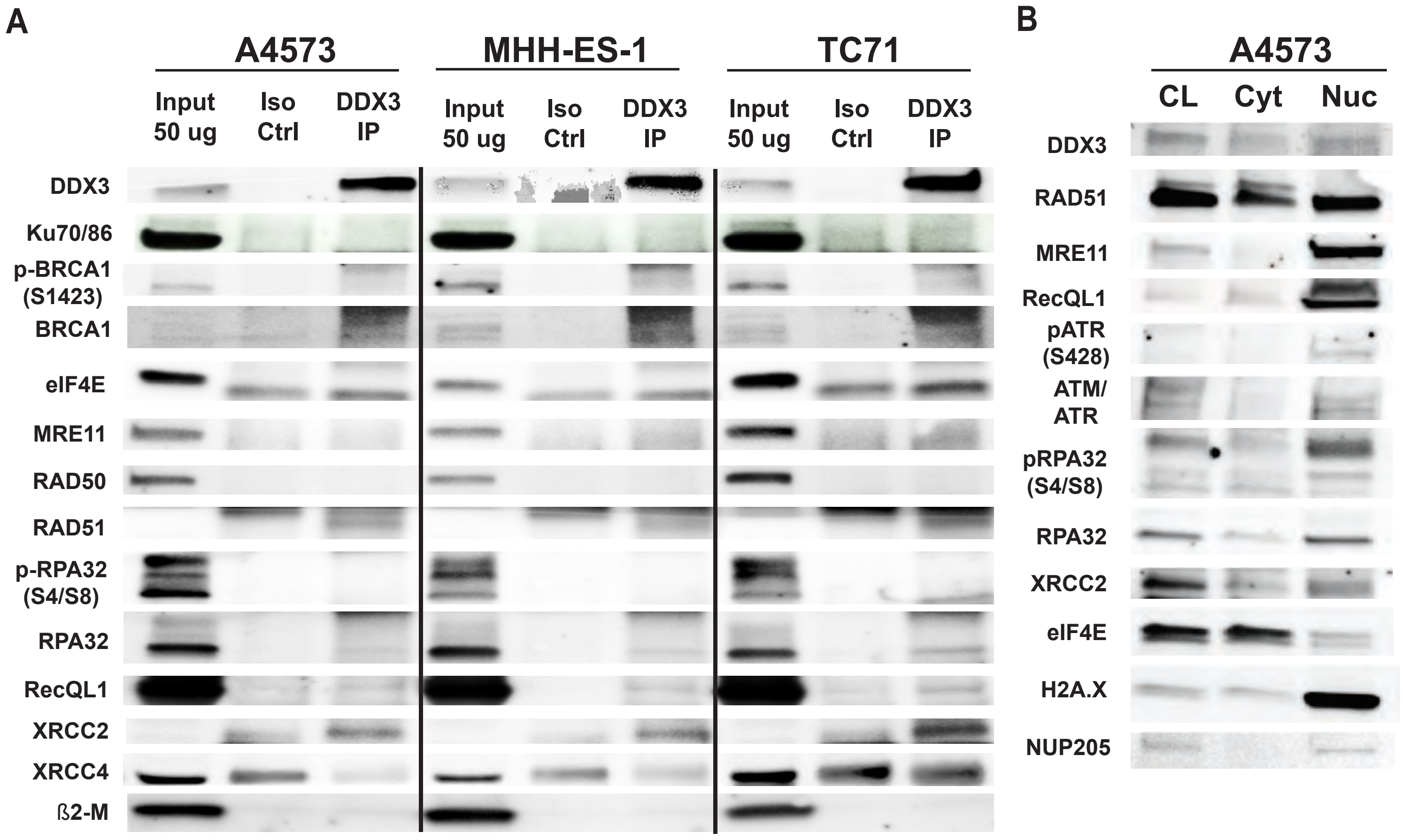

### Supplemental Figure 3

Supplemental Figure 3

#### Supp Figure 3

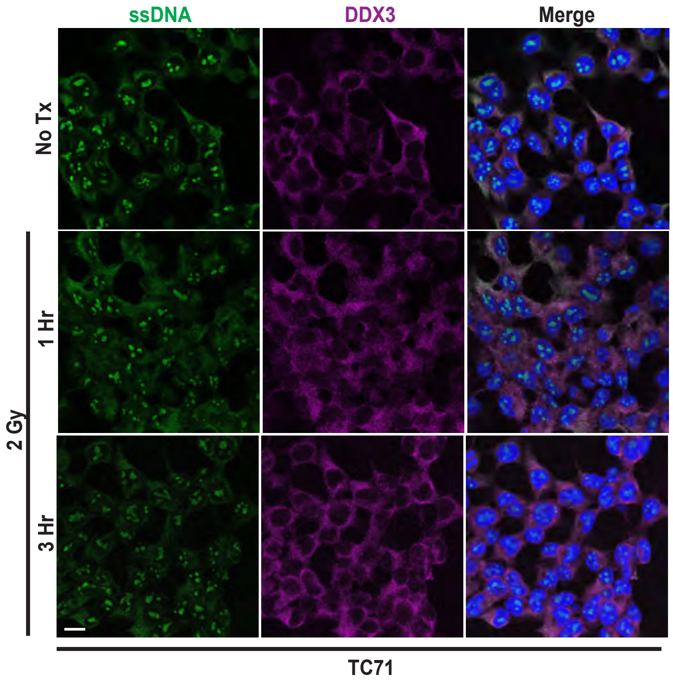
