## Supplemental Table 1 for "RNA Helicase DDX3 Regulates RAD51 Localization and DNA Damage Repair in Ewing Sarcoma"

| Primary Antibody | Molecular Weight | Isotype | Species | Application | Dilution/ Concentration | Cat. # | Company | RRID # |
|---|---|---|---|---|---|---|---|---|
| ATM | 350 | IgG | Rabbit | WB | 1:1000 | 2873 | Cell Signaling | AB_2062659 |
| ATR | 300 | polyIgG | Rabbit | WB | 1:1000 | 2790 | Cell Signaling | AB_2227860 |
| Beta-actin | 45 | IgG | Rabbit | WB | 1:10,000 | 4970 | Cell Signaling | AB_2223172 |
| Beta 2-microglobulin | 12 | IgG2b k | Mouse | WB | 1:5000 | sc-13565 | Santa Cruz Biotech. | AB_626748 |
| Beta 2-microglobulin | 12 | IgG | Rabbit | WB | 1:5000 | 12851 | Cell Signaling | AB_2716551 |
| BRCA1 | 220 | polyIgG | Rabbit | WB | 1:1000 | 9010 | Cell Signaling | AB_2228244 |
| Caspase 3 | 17, 32 | polyIgG | Rabbit | WB | 1:500 | ab13847 | abcam | AB_443014 |
| Chk1 | 56 | IgG2a k | Mouse | WB | 1:1000 | sc-8408 | Santa Cruz Biotech. | AB_627257 |
| Chk1 | 56 | IgG | Rabbit | WB | 1:1000 | ab47574 | abcam | AB_869133 |
| DDX3 (aa 1-114) Clone C-4 | 73 | IgG2b k | Mouse | WB; IF | 1:1000; 1μg/ml | sc-365768 | Santa Cruz Biotech. | AB_10844621 |
| DDX3 (aa 1-662) | 73 | IgG | Rabbit | WB | 1:1000 | ab235940 | abcam | AB _2910140 |
| DDX3 (aa 1-142) Clone AO196 | 73 | IgG1 k | Mouse | IHC | 1:50 | MABE1921-25UL | Sigma Aldrich | AB_2936197 |
| eIF4E | 28 | IgG1 k | Mouse | WB | 1:1000 | sc-9976 | Santa Cruz Biotech. | AB_627502 |
| H2A.X | 15 | IgG | Rabbit | WB | 1:1000 | 2595 | Cell Signaling | AB_10694556 |
| HA tag | | IgG1 | Mouse | IF | 2 μg/ml | H9658 | Sigma-Aldrich | AB_260092 |
| J2 | | IgG2a k | Mouse | IF | 2 μg/ml | 76651L | Cell Signaling | AB_2936194 |
| Ku70 | 70 | IgG2b k | Mouse | WB | 1:1000 | sc-71469 | Santa Cruz Biotech. | AB_1125206 |
| Ku86 | 86 | IgG1 | Mouse | WB | 1:1000 | sc-5280 | Santa Cruz Biotech. | AB_672929 |
| Mre11 | 79 | IgG1 | Mouse | WB | 1:1000 | ab214 | abcam | AB_302859 |
| NUP205 | 205 | IgG | Rabbit | WB | 1:1000 | PA5-55112 | ThermoFisher Scientific | AB_2644895 |
| p-ATR (S428) | 300 | polyIgG | Rabbit | WB | 1:1000 | 2853 | Cell Signaling | AB_2290281 |
| p-BRCA1 (S1423) | 220 | polyIgG | Rabbit | WB | 1:1000 | ab194753 | abcam | AB _2910141 |
| p-Chk1 (S345) | 56 | IgG | Rabbit | WB | 1:1000 | 2348 | Cell Signaling | AB_331212 |
| p-H2A.X (S139) | 15 | IgG1 | Mouse | IF | 2 μg/ml | 05-636 | Millipore Sigma | AB_309864 |
| p-H2A.X (S139) | 15 | IgG | Rabbit | WB | 1:1000 | 9718 | Cell Signaling | AB_2118009 |
| p-RPA32 (S4/S8) | 32 | polyIgG | Rabbit | WB | 1:2500 | A300-245A | Bethyl Laboratories | AB_210547 |
| Rad50 | 153 | IgG1 | Mouse | WB | 1:1000 | ab89 | abcam | AB_2176935 |
| Rad51 | 37 | IgG1 | Mouse | WB | 1:500 | sc-377467 | Santa Cruz Biotech. | AB _2910142 |
| Rad51 | 37 | IgG | Rabbit | IF | 2 μg/ml | ab63801 | abcam | AB _1142428 |
| Rad52 | 48 | IgG | Rabbit | WB | 1:1000 | ab124971 | abcam | AB_10971685 |
| RecQL1 | 75 | IgG2a k | Mouse | WB | 1:2500 | sc-166388 | Santa Cruz Biotech. | AB _2178425 |
| RPA32 | 32 | IgG1 | Mouse | WB | 1:1000 | ab2175 | abcam | AB _302873 |
| RPA32 | 32 | IgG | Rabbit | WB | 1:1000 | ab76420 | abcam | AB _1524336 |
| S9.6 (RNA:DNA hybrids) | | IgG2a | Mouse | IF | 75 μg/ml | gift | Dr. Teresa Bowman | N/A |
| S9.6 (RNA:DNA hybrids) | | IgG | Rabbit | IF | 2 μg/ml | Kf-Ab01137-23.0 | Kerafast | AB_2936195 |
| ssDNA | | IgG3 | Mouse | IF | 67 μg/ml | MAB3868 | EMD Millipore | AB _570342 |
| XPG | 200 | IgG | Rabbit | WB | 1:1000 | 11331-1-AP | Proteintech | AB_2098155 |

| | | | | | | | | |
|---|---|---|---|---|---|---|---|---|
| XRCC2 | 34 | IgG2a k | Mouse | WB | 1:500 | sc-365854 | Santa Cruz Biotech. | AB_10846464 |
| XRCC4 | 55 | IgG2a k | Mouse | WB | 1:500 | sc-271087 | Santa Cruz Biotech. | AB_10612396 |

| **Isotype Antibody** | **Species** | **Application** | **Concentration** | **Cat. #** | **Company** | **RRID #** |
|---|---|---|---|---|---|---|
| IgG1 | Mouse | IF | 2 µg/ml | MA1-10407 | ThermoFisher | AB_ 2536775 |
| IgG2a k | Mouse | IF | 40 µg/ml | 14-4724-85 | eBioscience | AB_470115 |
| IgG2b k | Mouse | IF | 1 µg/ml | 02-6300 | ThermoFisher | AB_2532949 |
| IgG3 | Mouse | IF | 2 µg/ml | 14-4724-82 | Invitrogen | AB_ 470120 |
| IgG | Rabbit | IF | 2 µg/ml | MA5-16384 | ThermoFisher | AB_ 2537903 |

| **Secondary Antibodies** | **Conjugate** | **Species** | **Application** | **Dilution/ Concentration** | **Catalogue #** | **Company** | **RRID #** |
|---|---|---|---|---|---|---|---|
| α-mouse IgG | HRP | Horse | WB | 1:10,000 | PI-2000 | Vector Laboratories | AB_ 2336177 |
| α-mouse IgG1 | AF488 | Goat | IF | 8 µg/ml | A21121 | Invitrogen | AB_ 2535764 |
| α-mouse IgG1 | AF555 | Goat | IF | 8 µg/ml | A21127 | Invitrogen | AB_ 2535769 |
| α-mouse IgG2a k | AF488 | Goat | IF | 8 µg/ml | A21131 | Invitrogen | AB_ 2535771 |
| α-mouse IgG2a k | AF594 | Goat | IF | 8 µg/ml | A21135 | Invitrogen | AB_ 2535774 |
| α-mouse IgG2a k | AF647 | Goat | IF | 8 µg/ml | A21241 | Invitrogen | AB_ 2535810 |
| α-mouse IgG2b k | AF594 | Goat | IF | 8 µg/ml | A21145 | Invitrogen | AB_ 2535781 |
| α-mouse IgG2b k | AF647 | Goat | IF | 8 µg/ml | A21242 | Invitrogen | AB_ 2535811 |
| α-mouse IgG3 | AF488 | Goat | IF | 8 µg/ml | A21151 | Invitrogen | AB_ 2535784 |
| α-rabbit IgG | AF555 | Goat | IF | 8 µg/ml | A21428 | Invitrogen | AB_ 2535849 |
| α-rabbit IgG | HRP | Goat | WB | 1:10,000 | PI-1000 | Vector Laboratories | AB_ 2336198 |
